# Historical population declines prompted significant genomic erosion in the northern and southern white rhinoceros (*Ceratotherium simum*)

**DOI:** 10.1101/2020.05.10.086686

**Authors:** Fátima Sánchez-Barreiro, Shyam Gopalakrishnan, Jazmín Ramos-Madrigal, Michael V. Westbury, Marc de Manuel, Ashot Margaryan, Marta M. Ciucani, Filipe G. Vieira, Yannis Patramanis, Daniela C. Kalthoff, Zena Timmons, Thomas Sicheritz-Pontén, Love Dalén, Oliver A. Ryder, Guojie Zhang, Tomás Marquès-Bonet, Yoshan Moodley, M. Thomas P. Gilbert

**Affiliations:** GLOBE Institute, University of Copenhagen, Øster Voldgade 5-7, 1350 Copenhagen, Denmark; Institut de Biologia Evolutiva (Consejo Superior de Investigaciones Científicas–Universitat Pompeu Fabra), Barcelona Biomedical Research Park, Doctor Aiguader 88, 08003 Barcelona, Catalonia, Spain; Department of Zoology, Swedish Museum of Natural History, Frescativägen 40, 114 18 Stockholm, Sweden; Department of Natural Sciences, National Museums Scotland, Chambers Street, Edinburgh, EH1 1JF, Scotland, United Kingdom; Centre for Palaeogenetics, Svante Arrhenius väg 20C, 10691 Stockholm, Sweden; Department of Bioinformatics and Genetics, Swedish Museum of Natural History, Frescativägen 40, 114 18 Stockholm, Sweden; San Diego Zoo Institute for Conservation Research, 15600 San Pasqual Valley Road, Escondido, 92027 CA, USA; Section for Ecology and Evolution, Department of Biology, University of Copenhagen, 2100 Copenhagen, Denmark; State Key Laboratory of Genetic Resources and Evolution, Kunming Institute of Zoology, Chinese Academy of Sciences, 650223, Kunming, China; Center for Excellence in Animal Evolution and Genetics, Chinese Academy of Sciences, 650223, Kunming, China; BGI-Shenzhen, 518083, Shenzhen, China; Department of Zoology, University of Venda, University Road, 0950 Thohoyandou, Republic of South Africa; Norwegian University of Science and Technology, University Museum, 7491 Trondheim, Norway; National Centre for Genomic Analysis–Centre for Genomic Regulation, Barcelona Institute of Science and Technology, 08028 Barcelona, Spain; Institucio Catalana de Recerca i Estudis Avançats (ICREA), 08010 Barcelona, Catalonia, Spain; DTU Bioinformatics, Kongens Lyngby, Hovedstaden 2800, Denmark; Centre of Excellence for Omics-Driven Computational Biodiscovery (COMBio), Faculty of Applied Sciences, AIMST University, Kedah, Malaysia

**Keywords:** northern white rhinoceros, southern white rhinoceros, population decline, genomic erosion, delta estimators, conservation genomics

## Abstract

Large vertebrates are extremely sensitive to anthropogenic pressure, and their populations are declining fast. The white rhinoceros (*Ceratotherium simum*) is a paradigmatic case: this African megaherbivore suffered a remarkable population reduction in the last 150 years due to human activities. The two white rhinoceros subspecies, the northern (NWR) and the southern white rhinoceros (SWR), however, underwent opposite fates: the NWR vanished quickly after the onset of the decline, while the SWR recovered after a severe bottleneck. Such demographic events are predicted to have an erosive effect at the genomic level, in connection with the extirpation of diversity, and increased genetic drift and inbreeding. However there is currently little empirical data available that allows us to directly reconstruct the subtleties of such processes in light of distinct demographic histories. Therefore to assess these effects, we generated a whole-genome, temporal dataset consisting of 52 re-sequenced white rhinoceros genomes, that represents both subspecies at two time windows: before and during/after the bottleneck. Our data not only reveals previously unknown population substructure within both subspecies, but allowed us to quantify the genomic erosion undergone by both, with post-bottleneck white rhinoceroses harbouring significantly fewer heterozygous sites, and showing higher inbreeding coefficients than pre-bottleneck individuals. Moreover, the effective population size suffered a decrease of two and three orders of magnitude in the NWR and SWR respectively, due to the recent bottleneck. Our data therefore provides much needed empirical support for theoretical predictions about the genomic consequences of shrinking populations, information that is relevant for understanding the process of population extinction. Furthermore, our findings have the potential to inform management approaches for the conservation of the remaining white rhinoceroses.

## Introduction

Earth’s biodiversity is experiencing a severe crisis. In the past 100 years, species have gone extinct at rates that are up to 100-fold higher than conservative estimates of background extinction rates [1]. But deterioration of biodiversity is also pervasively manifested by shrinking population sizes, population extirpations and fragmentation of ranges of distribution in extant species [2]. Alarmingly, these pervasive threats to biodiversity can be tightly linked to anthropogenic activities [1,3,4].

Large vertebrates are particularly sensitive to the impact of anthropogenic pressures, and their wild populations have undergone marked declines [4]. Beyond the net loss of individuals and populations, these rapid declines can potentially have dire genetic consequences [5]. Theory predicts that populations undergoing dramatic and rapid decreases in size (i.e. bottlenecks), will also lose substantial genetic diversity and subsequently suffer from strong genetic drift and inbreeding [6,7]. All this, in turn, can potentially decrease fitness [8], and in the long term, diminish resilience to environmental change [6].

In an era where genomic-scale data generation has become increasingly tractable for non-model species, and ancient DNA techniques allow us to retrieve such information from even long-dead biological material, the opportunity now exists to directly observe signals of the demographic history of wild populations [9–13]. In the case of declining populations, this implies being able to detect and measure *genomic erosion* as the suite of symptoms characteristic of such demographic trajectories, mainly lower levels of genomic diversity and heterozygosity [13].

The white rhinoceros (*Ceratotherium simum* Burchell) is an exemplary case of the fate of the megafauna in the Anthropocene. This species is divided into two allopatric populations, also considered subspecies: the northern white rhinoceros (*Ceratotherium simum cottoni*, NWR in this text), and the southern white rhinoceros (*Ceratotherium simum simum*, SWR in this text) [14]. Because of their obligate grazing lifestyles, both are bound to sub-Saharan African grasslands, but their geographical ranges are non-overlapping [15,16], they are genetically distinct [14,17,18], and they have not come into genetic contact for at least 10,000 years [18]. They have undergone remarkably different demographic trajectories in the past 150 years that share, however, one striking similarity: the occurrence of a dramatic, human-driven population decline [15,16,19].

The SWR roamed south of the Zambezi river across present-day South Africa, Botswana, Namibia, Zimbabwe and Mozambique (Figure 1A), until the expansion of European colonialism in southern Africa in the 19^th^ century. Habitat clearance and hunting pushed the SWR through a bottleneck from an estimated size of several hundred thousands, to an estimated low of 200 individuals at the turn of the 20^th^ century [19]. Conservation efforts, however, boosted a remarkable recovery of the only surviving population in Kwa-Zulu (South Africa) [19]. There are currently ∼18,000 wild individuals [20], although they remain threatened due to the relentless poaching for their highly valued horn.

**Figure 1.**
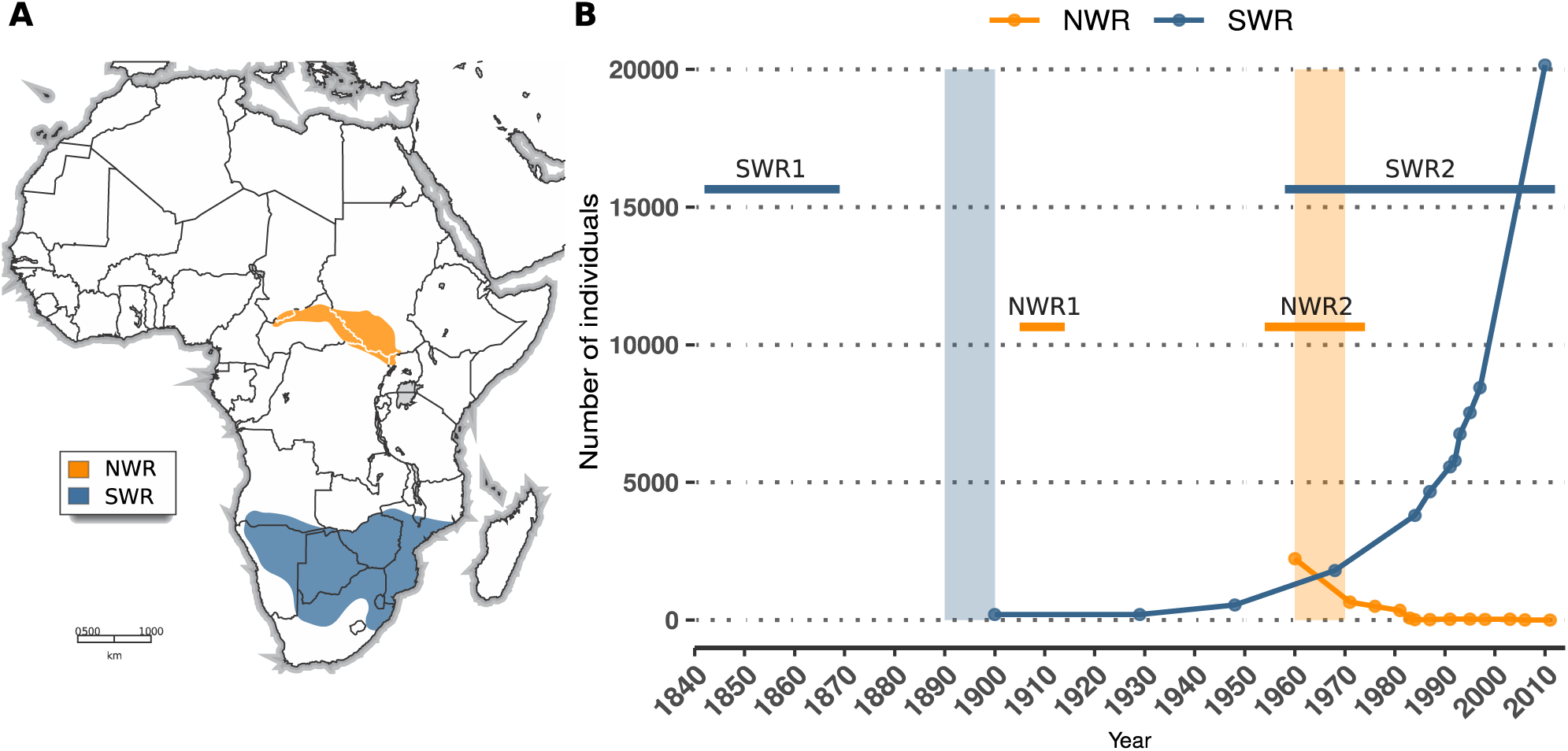
Ranges of distribution and recent demography of the northern and southern white rhinoceros. A) The historical distribution of the NWR (*C. s. cottoni*) and the SWR (*C. s. simum*) (map adapted from [15]). B) Recent demographic histories of the NWR and the SWR according to census size estimates reported in the literature [15,19,20]. Absolute values prior to these dates do not exist, but population sizes are known to have been larger from historical records [15,19,20]. Vertical shades indicate approximate timing of bottlenecks. Horizontal bars indicate our sampling time windows, which in NWR1 and SWR1 refer to dates of collection, and in NWR2 and SWR2, to dates of birth.

In contrast, the NWR did not recover from the poaching onslaught that began in the mid-20^th^ century. Once inhabiting the plains of present-day South Sudan, Northeastern Democratic Republic of the Congo (DRC), Central African Republic and Uganda (Figure 1A), the NWR was rapidly decimated in subsequent poaching bursts boosted by civil and political instability in the area [15,16,21]. Although in 1960 there were still ∼2000 NWR in the wild, by 1984, only 350 remained [15]. The NWR was declared extinct in the wild in 2011 by the IUCN [20].

A genetic study revealed a loss of diversity at mitochondrial and microsatellite markers, as a consequence of these demographic histories, when comparing individuals born during or after the population bottleneck, against older samples collected from museums [17]. To extend such analyses to the genome level, and test whether either subspecies have undergone genomic erosion or other effects as a result of their recent demographic history, we generated a re-sequenced, whole-genome dataset for both NWR and SWR populations, sampled at two time windows each throughout the past ∼170 years. With this dataset, we investigated a) the population structure within and among these pre- and post-bottleneck white rhinoceroses, b) the occurrence and magnitude of genomic erosion, and c) the effect of the recent demographic trajectories on the effective population size (N_e_) of the NWR and the SWR.

## Results and discussion

We generated the most comprehensive white rhinoceros genomic dataset to date, consisting of 52 genomes re-sequenced at an average depth of coverage of 12.4× (see Table S2 and Figure S2 for further details). Of these, 30 were newly generated genomes from historical museum specimens, and 22 were modern genomes, of which 13 were publicly available (see [18]) and 9 were newly re-sequenced (Table S1). These 52 samples represent both NWR (n = 25) and SWR (n = 27) from different geographical locations, at two time windows each (Figure 1A), hereafter referred to as pre- and post-bottleneck, with the suffix 1 (e.g. NWR1) for the former, and 2 (e.g. NWR2) for the latter.

Sample names specify first the alpha-2 code of the country of origin, followed by the year of collection for historical samples, and year of birth for modern individuals; a subsequent index number distinguishes samples from the same country and year. Among post-bottleneck samples, all NWR2 were wild-born animals taken into captivity, while within SWR2, some were wild (n = 12) and some were bred in captivity (n = 6) individuals, which is denoted by the prefix ‘cap’ in the sample name.

Demographic histories and sampling windows differ between subspecies. For the NWR, a time-span of ∼70 years is covered, during which a population decline was followed by a rapid collapse (Figure 1B). The SWR, on the other hand, are represented in our study by samples from two time windows at the ends of a period of ∼170 years, during which a strong bottleneck was followed by a recovery (Figure 1B).

### Geography and potential sampling gaps drive the structure in the dataset

We ran a principal component analysis (PCA) on all unrelated individuals in the dataset (n = 49) (see *Relatedness test* and Figure S3 in Supplementary material), that revealed a clear separation between NWR and SWR (Figure S4), and suggested the existence of substructure within each subspecies. To further resolve the fine patterns of structure, we conducted a non-linear dimensionality reduction analysis with UMAP [22]. With this approach, we identified previously unknown historical population substructure in the NWR and the SWR, seemingly driven by geography (Figure 2A).

**Figure 2.**
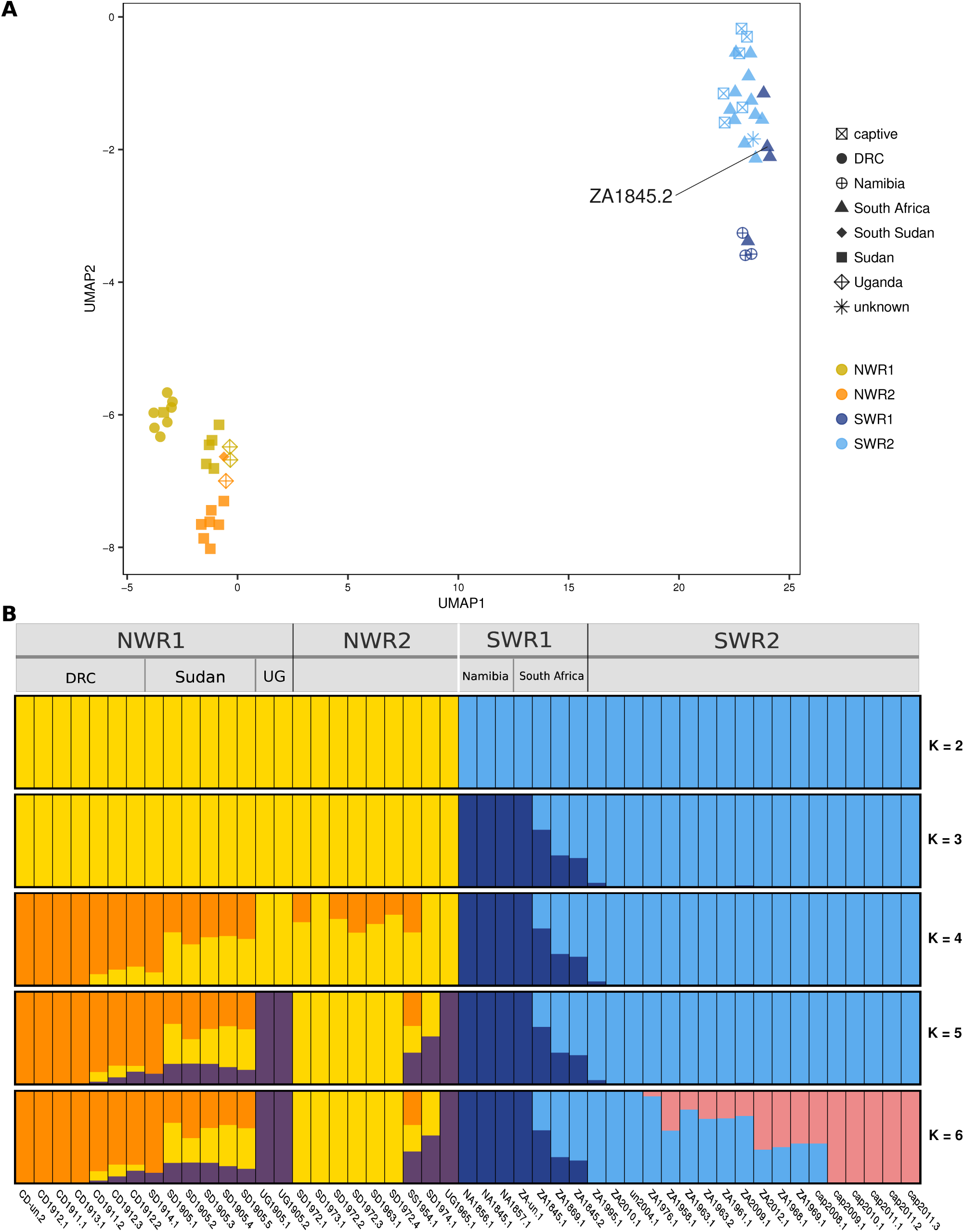
The structure of genomic variation in northern and southern white rhinoceros. A) Visualization of the two axes of the UMAP manifold in an analysis of 49 unrelated individuals; the labelled SWR1 sample (ZA1845.2) corresponds to a historical individual known to have been sourced from Kwa-Zulu. C) Admixture analysis of 49 unrelated individuals. The run of highest likelihood of a total of 50 runs is displayed for each value of K.

In the NWR, pre-bottleneck individuals featured two clusters corresponding to current Sudan-Uganda and the Democratic Republic of the Congo (DRC) (Figure 2A). This may be a true biological signal of some degree of isolation between the two groups, but we caveat that it could be an artefact of insufficient sampling across a continuous cline of diversity. All post-bottleneck NWR diversity in our dataset fell among or close to the Sudan-Uganda historical diversity, in agreement with the zoo studbooks that corroborate that those were the countries of origin of these imported wild animals.

In the SWR, pre-bottleneck samples also showed structure, with Namibian and one South African samples forming a first group, while three other South African samples constituted a second group to which all post-bottleneck SWR were closely related (Figure 2A). Metadata regarding the location of origin of most SWR1 samples was scarce, however ZA1845.2 was known to have been collected in Kwa-Zulu (South Africa). This sample also clustered closely with two other historic SWR samples (Figures 2A and S4), suggesting that they too could be from the same or adjacent geographical region. Kwa-Zulu was the only SWR population that survived the bottleneck of the late 19^th^ century, and was the source of all modern SWR. This was supported by our UMAP analysis (Figure 2A), which showed all modern SWR samples clustering with ZA1845.2.

Admixture analysis of 49 unrelated individuals, with values of ancestral components (K) ranging between two and six, showed concordant trends. Results featured a clear separation between NWR and SWR at K = 2 (Figure 2B), while subsequent values of K separated subclusters following geographical patterns. Among the NWR, DRC separated first from the Sudan-Uganda samples, and these in turn split next (Figure 2B). As in the UMAP analysis, post-bottleneck NWR contain ancestry mainly from Sudan and Uganda. Pre-bottleneck SWR display also some local substructure, with a western group (three Namibian and one South African samples), and an eastern group of South African individuals, possibly from Kwa-Zulu (it includes ZA1845.2). Post-bottleneck SWR are all contained within the latter eastern group (Figure 2B).

Overall, the data suggests that there was substructure in historical times within both the NWR and the SWR, driven principally by geography. Given the potential gaps in our sampling scheme, and the absence of obvious barriers to gene flow, the clusters we observed within each subspecies, however, might have been part of a more continuous cline caused by isolation by distance over an uninterrupted distribution. Additionally, the founder effect that gave rise to all extant SWR (caused by population extirpation) is detectable in our dataset.

### Post-bottleneck white rhinoceroses show signs of genomic erosion

To quantify genomic erosion within each subspecies, we computed measures of genomic diversity per group and per individual, and then calculated the difference between time windows as the proportional difference between the post- and pre-bottleneck median value of each metric. These intra-species temporal measures of genomic change represent *delta estimators* as first proposed in [13].

We first estimated π_tv_ as a proxy for nucleotide diversity (average number of pairwise differences across a set of sequences), including only transversions so as to conservatively exclude any possible overinflation driven by post mortem cytosine deaminations [23,24]. The group median of π_tv_ across 63 scaffolds was 5.22% lower among NWR2 than in NWR1 (paired samples Wilcoxon test p-value = 6.4 × 10^−4^), and 27.67% lower in SWR2 compared to SWR1 (paired samples Wilcoxon test p-value = 5.3 × 10^−12^) (Figure 3A, Table 1).

**Table 1.** D*elta estimators for measures of genomic diversity between pre- and post-bottleneck white rhinoceroses*. Median values of π_tv_, genome-wide heterozygosity (GW het), heterozygosity at exons (exon het) and F_roH_ for each of the four groups. Comparisons of the medians between NWR1 and NWR2, and between SWR1 and SWR2 were calculated with Wilcoxon tests. Delta estimators were calculated as median pre-bottleneck value minus median post-bottleneck value, divided by the median pre-bottleneck value. All metrics were based on transversion sites only.

| | n | $\pi_{tv}$ | Paired<br>Wilcoxon<br>test p-value | $\Delta\pi_{tv}$ | Median<br>GW het | Unpaired<br>Wilcoxon<br>test p-value | $\Delta$<br>GW<br>het | Median<br>exon het | Unpaired<br>Wilcoxon<br>test p-value | $\Delta$<br>exon<br>het | Median<br>$F_{RoH}$ | Unpaired<br>Wilcoxon<br>test p-value | $\Delta$<br>$F_{RoH}$ |
| --- | --- | --- | --- | --- | --- | --- | --- | --- | --- | --- | --- | --- | --- |
| NWR1 | 16 | 0.000345 | 6.40E-04 | -0.0522 | 0.000202 | 1.96E-06 | -0.1284 | 0.000117 | 1.07E-03 | -0.1785 | 0.1679 | 3.28E-02 | 0.2710 |
| NWR2 | 9 | 0.000327 |  |  | 0.000176 |  |  | 0.000096 |  |  | 0.2134 |  |  |
| SWR1 | 9 | 0.000253 | 5.30E-12 | -0.2767 | 0.000158 | 1.28E-05 | -0.3549 | 0.000087 | 2.36E-03 | -0.3194 | 0.1831 | 1.68E-03 | 0.7112 |
| SWR2 | 18 | 0.000183 |  |  | 0.000102 |  |  | 0.000060 |  |  | 0.3133 |  |  |

**Figure 3.**
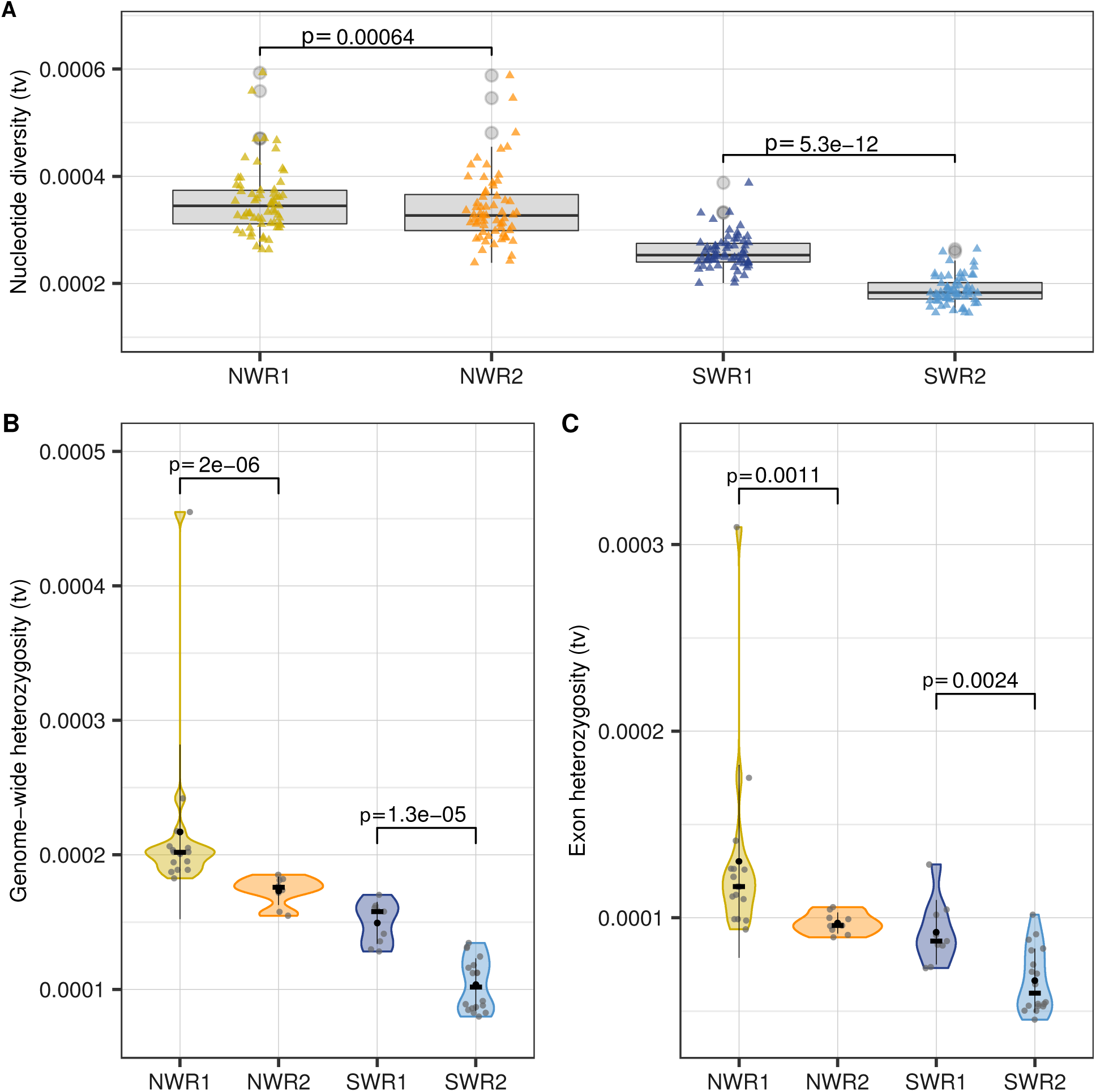
Post-bottleneck white rhinoceroses show lower genomic diversity. A) Distribution of values of π_tv_ across 63 scaffolds for each of the four groups. Each coloured triangle represents the value of π_tv_ for one scaffold. P-values correspond to comparisons of medians with paired Wilcoxon tests. B) Estimates of individual genome-wide heterozygosity, and (C) heterozygosity at regions annotated as exons, based on the per-sample SFS calculated for transversions and then corrected for depth of coverage. Black dots indicate the mean, black lines the standard deviation, and black cross-bars, the median per group. P-values correspond to median comparisons with unpaired Wilcoxon tests.

The individual genome-wide heterozygosity (corrected for depth of coverage; see *Heterozygosity correction* in Methods and Figure S5) was also lower among post-bottleneck individuals (Figure 3B). Median values between NWR1 and NWR2, and between SWR1 and SWR2 were significantly different (unpaired samples Wilcoxon test p-value_NWR_ = 1.96 × 10^−6^; unpaired samples Wilcoxon test p-value_SWR_ = 1.28 × 10^−5^). In fact, NWR2 showed 12.84% lower heterozygosity than NWR1, and SWR2 featured a median heterozygosity that is 35.49% lower than among SWR1 (Table 1).

These significant losses in genomic diversity are signs of genomic erosion that might be affecting the genome indiscriminately. To verify this, we investigated specifically whether coding regions might have selectively retained pre-bottleneck levels of diversity instead, given their functional importance. We calculated an individual measure of heterozygosity across regions annotated as exons in the white rhinoceros reference assembly, and calculated the delta estimators between time windows. As a result we detected a decline between pre- and post-bottleneck groups comparable to that of genome-wide heterozygosity (Figure 3C). The NWR2 exhibited a decay of 17.85% in exon heterozygosity compared to NWR1 (unpaired samples Wilcoxon test p-value_NWR_ = 1.07 × 10^−3^); SWR2 featured a decrease of 31.94% (unpaired samples Wilcoxon test p-value_SWR_ = 2.36 × 10^−3^) (Table 1).

This reflects a pervasive loss of diversity along the genome, even in regions of high functional importance whose erosion might trigger detrimental phenotypic effects. Although strong negative selection, i.e. purging, could have acted to retain diversity in exons, the time span considered might have been too short for highly deleterious phenotypic effects to arise, as well as for balancing selection to preserve diversity.

Given the well documented sharp decline in population sizes for both NWR and SWR since the 19^th^ century [15], we hypothesised that signs of recent inbreeding might have accumulated in their genomes [13]. We therefore estimated inbreeding per sample as the fraction of the genome putatively in runs of homozygosity (RoH), according to local estimates of heterozygosity in sliding windows of 1 Mbp (0.5 Mbp slide) without further correction.

Based on the distribution of estimates of heterozygosity per sample (Figure S7), we set a threshold of heterozygosity at 5 × 10^−5^ below which two or more consecutive windows harbouring such small values are considered to constitute a RoH. We excluded from RoH assessments the three samples of lowest depth of coverage (CD-un.1, CD1911.2 in NWR1, and ZA1845.1 in SWR1), since they displayed an excess of local heterozygosity estimates equal or close to zero due to missing data (Figure S7).

We first observed that post-bottleneck individuals harboured the longest detectable RoHs, particularly SWR2 (Figure 4A). We then computed F_RoH_ per sample as the ratio between the length (in base pairs) falling within RoH and the total length scanned. Median F_RoH_ values were significantly higher among post-bottleneck samples in both subspecies (unpaired samples Wilcoxon test p-value_NWR_ = 3.28 × 10^−2^; unpaired samples Wilcoxon test p-value_SWR_ = 1.68 × 10^−3^) (Figure 4B). The NWR experienced an increase of F_RoH_ of 27.10%, while the difference rose to 71.12% in the SWR (Table 1).

**Figure 4.**
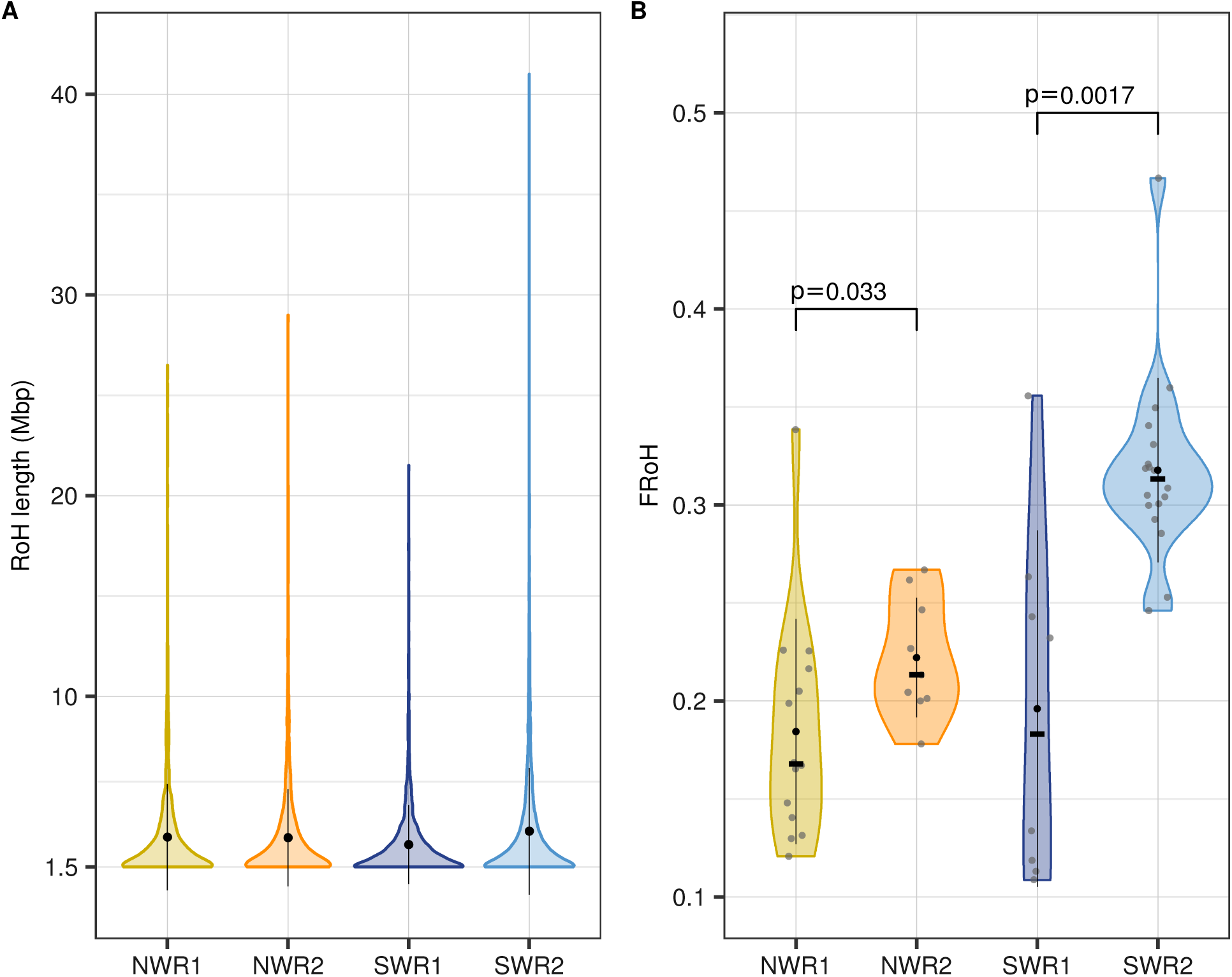
Post-bottleneck white rhinoceroses show higher estimates of inbreeding. A) Distribution of the length of RoHs for each of the four groups; black dots indicate the mean RoH length per group and black lines the standard deviation. B) Estimates of individual F_RoH_ across groups. Black dots indicate the mean, black lines the standard deviation, and black cross-bars the median per group. P-values refer to unpaired Wilcoxon tests to compare the medians.

This contrast between subspecies in the magnitude of change of F_RoH_ might reflect the difference in time-span between sampling time windows, but also the disparity of their demographic histories. Specifically, the NWR was much more rapidly reduced than the SWR, thus the SWR simply had more time to accumulate symptoms of inbreeding in their genomes.

We caution, however, that the local estimates of heterozygosity on which we based our RoH analyses were not corrected for depth of coverage, so variability in depth among samples (despite the removal of three outliers) might still have an effect. Delta estimators for F_RoH_ must be therefore interpreted carefully. Additionally, the available reference genome of the white rhinoceros is yet to be assembled at chromosome level, so identifying RoH with precision is not feasible. Even if coarse, however, this approach provides a first approximation to estimating inbreeding with genomic data when robust population allele frequencies are not available.

To exclude the fact that sampling biases might be creating spurious patterns of genomic erosion, we calculated the individual metrics of genomic diversity for a subset of the samples. For NWR1, we retained only samples originating from Sudan-Uganda, since all NWR2 were sourced from that area. And for SWR2, only wild-born individuals were retained for comparison with all SWR1. This subsampled dataset consisted of 38 individuals, distributed across the four groups (see Table S3), among which we still detected a significant loss of diversity between post- and pre-bottleneck time windows.

Both Sudan-Uganda NWR2 and wild SWR2 showed significantly lower values of genome-wide and exon heterozygosity compared to their pre-bottleneck counterparts (Figure S8), and the distribution of RoH length changed only slightly in this subsampled dataset (Figure S9A). Delta estimators displayed comparable values to those for the entire dataset (Table S3), with the exception of F_RoH_ in the NWR, which was not significantly different between pre- and post-bottleneck Sudan-Uganda NWR (Figure S9B and Table S3). Given their rapid disappearance, it is plausible that they did not suffer strong inbreeding over several generations, but the bottleneck itself probably caused the observed loss of genomic diversity. In the SWR, the intensity of the erosion remains significant and higher across all metrics (Table S3), probably owing to the occurrence of both population extirpation during the bottleneck, and subsequent genetic drift and inbreeding in the surviving wild SWR.

### Their recent demographic histories have reduced N_e_ in the NWR and the SWR

The effective size of a population (N_e_) is the size of an idealised Wright-Fisher population that loses genetic diversity at the same rate as the real population, and can be interpreted as an estimator of the variability available in a population at a given time to source the next generation [6]. The census size and genomic diversity of the NWR and the SWR decreased between the two sampling windows in our dataset. Thus we assessed whether this triggered substantial changes in N_e_ by modelling the recent demographic histories in a coalescent framework with fastsimcoal2 [25].

Our simulations reflected the known demographic trajectories of SWR and NWR from historical records. For the NWR, we simulated a population bottleneck scenario without any further size change; whereas the SWR model included the same initial population bottleneck for some time, and then a subsequent population expansion. In each case we used the 2-dimensional SFS of the pre- and post-bottleneck samples for each subspecies, and we used a mutation rate of 2.5 × 10^−8^ mutations per generation as in [18].

The best-fitting parameter values indicated a decline of N_e_ of two (for NWR) and three (for SWR) orders of magnitude between the first sampling event and the bottleneck (Figure 5). In the SWR, the expansion after the bottleneck subsequently increased N_e_ again, so that at the second time window it grew to half the value in pre-bottleneck times (Figure 5).

**Figure 5.**
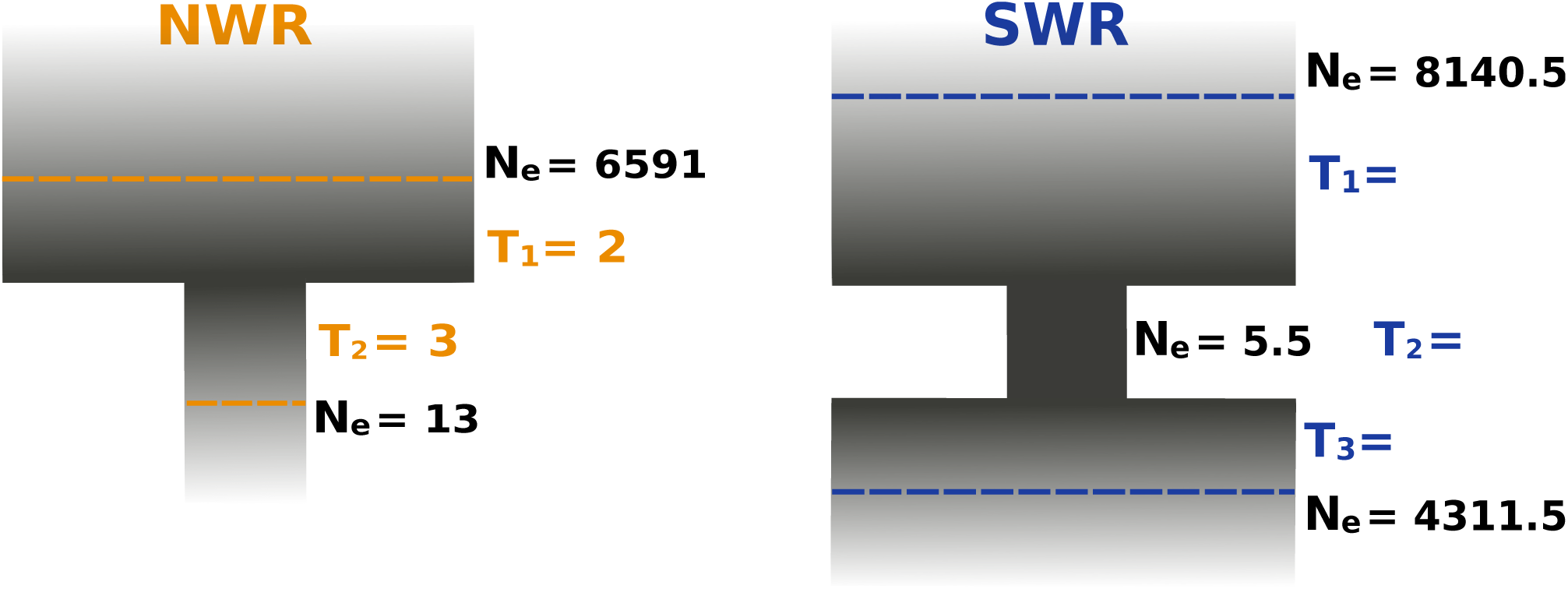
The recent demographic histories reduced N_e_ in the northern and southern white rhinoceros. Demographic models simulated with fastsimcoal2 [25] for the NWR and the SWR. Dashed lines indicate sampling time windows before and after the bottleneck. Best fitting values of N_e_ at each time window are expressed as number of diploid individuals. Estimates of time between events (first sampling, onset and end of the bottleneck and second sampling) are expressed in number of generations. Mutation rate of 2.5 × 10^−8^ mutations per generation [18].

We also used the simulations to estimate the time intervals, in generations, between events. For the NWR, the total ∼70 years between the first and the second sampling time windows were split into two intervals in the model: from the first sampling window to the bottleneck, and from there to the second sampling event. The first encompassed two generations according to the model, and the second represented three generations (Figure 5). In the SWR, the model estimated that the total length of ∼170 years between the first and the second sampling event corresponded to 26 generations, of which the bottleneck spanned 10 (Figure 5).

These time estimates would be congruent with a scenario in which the mean generation time of the species is ca. 8 years, as some authors have proposed [18]. However, this figure refers to the age at first birth of white rhinoceros females [26], and thus other authors have suggested that the median generation time for the species lies around 27 years [17]. Under this scenario, the numbers of generations between events in our models would be overestimates. This might be caused by the population substructure present in the pre-bottleneck groups, compared to the homogeneity of post-bottleneck individuals. Going backwards in time, many coalescence events are expected to happen at the time of the bottleneck; the remaining lineages to coalesce will take much longer to do so given that the pre-bottleneck populations had a higher N_e_ and were substructured.

Despite the potential overestimation of the number of generations between events, these models highlight the sizeable and relatively rapid impact of dramatic demographic events on the effective size of a population. The time in years covered by our dataset is relatively short, and yet remarkable oscillations of N_e_ were observed.

Just a century and a half ago the two subspecies of white rhinoceros were substructured populations distributed across two allopatric, broad ranges of distribution. Our results suggest that geography underlied this substructure. However, the uneven nature of museum-based sampling impedes concluding whether the clusters we observed were part of a continuous cline of diversity following a pattern of isolation by distance. Both subspecies suffered dramatic bottlenecks in their recent history, but their timing and outcomes differed: the NWR collapsed permanently in a few decades, while the SWR population size decreased but then re-expanded from a single surviving population (as detected in our structure analyses). Despite these opposite outcomes, the population declines left an imprint in both subspecies in the form of genomic erosion: heterozygosity levels decreased and inbreeding estimates increased.

The magnitude and specific profile of the genomic erosion was different between subspecies. In the NWR, for which both our sampling time windows, and the time until their disappearance, were closer in time than for the SWR, genomic erosion was less marked than in the SWR. We feel the shorter time scale of events in the NWR probably explains why the estimated inbreeding did not increase remarkably. The SWR, on the other hand, have been recovering from a strong founder effect, and suffering from drift and inbreeding until the present, which explains the larger magnitude of the genomic erosion they featured. Our results suggest that a careful control of inbreeding would be advisable in the management of the remaining SWR, given the notably high levels of F_RoH_ and RoH length when compared to historical counterparts.

As a reason for optimism, however, the demographic trajectory of the SWR in the last century proves that white rhinoceroses have the potential to reconstitute thriving populations from a very limited pool of genomic diversity. This is highly relevant as rewilding projects become more realistic. State-of-the-art techniques in molecular and reproductive biology, at the service of conservation, might be able to restore (or even increase) some of the rhinoceros genomic diversity lost in the wild [27,28].

Species extinctions are often used as a metric of anthropogenic pressure, but population extirpations and census size reductions are more pervasive, precede extinction, and affect both officially threatened and non-threatened species [2]. As a consequence, snippets of genomic diversity are being erased from the biosphere for good. The viability of a species might not be cancelled in the short-term as a result, but its genomic pool will be eroded, which is in itself a biodiversity loss, and potentially entails an added degree of vulnerability to the species in the face of a changing environment. Ultimately, if conservation of biodiversity is entering an era of tailor-made, population-specific approaches, temporal datasets hold great power to reveal the hidden consequences (and potential risks) of the human threat to biodiversity.

## Supporting information

Supplementary material (main)

## Acknowledgements

This work was supported by ERC Consolidator Grant 681396 ‘Extinction Genomics’ to M.T.P.G. and by EMBO Short-Term Fellowship 7578 to F.S.B. The authors would like to acknowledge support from Science for Life Laboratory, the National Genomics Infrastructure (NGI), Sweden, the Knut and Alice Wallenberg Foundation and UPPMAX for providing assistance in massively parallel DNA sequencing and computational infrastructure.

The authors are very grateful to all the museums who provided samples for this study: the American Museum of Natural History, the National Museums Scotland, the Natural History Museum at the National Museum Praha, the Natural History Museum Vienna, the Powell-Cotton Museum, the Royal Museum for Central Africa Tervuren, and the Swedish Museum of Natural History.

## Author Contributions

Conceptualization, F.S.B., S.G., J.R.M., M.V.W., M.M., F.G.V., Y.M. and M.T.P.G.; Investigation, F.S.B., M.C., L.D. and O.A.R.; Formal Analysis, F.S.B., S.G., J.R.M., M.V.W., M.M., A.M., and Y.P.; Data Curation, F.S.B., L.D. and O.A.R; Resources, D.C.K., Z.T., T.S.P., G.Z. and T.M.B.; Writing - Original Draft, F.S.B.; Writing –Review & Editing, F.S.B., S.G., J.R.M, M.V.W., M.M., M.C., F.G.V., D.C.K., Z.T., T.S.P., L.D., O.A.R., T.M.B, Y.M. and M.T.P.G; Visualization, F.S.B.;

Supervision, Y.M. and M.T.P.G.; Funding Acquisition, M.T.P.G

## Methods

### Data generation procedures

#### Processing of historical white rhinoceros material

Our historical sample collection included material obtained from 33 museum specimens. Collection dates ranged between 1845 and 2010. Most samples consisted of keratinous material as fragments or drill powder; a few, were bone fragments. Destructive sampling of all specimens took place at the institution of origin of the material. Samples from museum specimens were stored and processed in facilities dedicated to ancient DNA work at the Swedish Museum of Natural History, Stockholm, and the Natural History Museum of Denmark, Copenhagen.

Skin samples were manually cut with disposable scalpels in order to generate 20-80 mg of fragmented tissue. Dry skin tissue is highly absorbent, therefore the biggest pieces of material were hydrated for 2-3 hours at 4 °C by adding 0.5-1 mL of molecular biology grade water to each tube. Water was discarded, and the tissue was briefly washed with 0.5 mL of a 1% bleach solution, followed by two washes with molecular biology grade water. Bone material was crushed with a small hammer, and small pieces amounting to 150-200 mg were used for extraction.

#### DNA extraction, library preparation and sequencing

Extraction of DNA from keratinous tissue of museum specimens was done either with the DNeasy Blood and Tissue Kit (QIAGEN®) or with the method detailed in [29]. In the first case, two modifications were introduced to the manufacturer’s guidelines: adding of DTT (Dithiothreitol) 1M to a final concentration of 40 mM to the lysis buffer, and the substitution of the purification columns in the kit by MinElute® silica columns (QIAGEN®) to favour retention of small fragments. In the second approach, keratinous tissue was digested overnight at 56 °C, while rotating, in 800 Lof freshly made digestion buffer containing 10 mM Tris-HCl (pH 8.0), 10 mM NaCl, 2% SDS, 5 mM CaCl2, 2.5 mM EDTA (pH 8.0), 40 mM DTT, 1 µg/μL proteinase K solution and molecular biology grade water. Bone samples were digested overnight at 56 °C, while rotating, in 800 L of freshly made lysis buffer for bone tissue containing 0.4 mM EDTA (pH 8.0), 0.098 mM urea, and 0.22 µg/μL proteinase K.

Tubes were then centrifuged on a bench-top centrifuge at 6000 × *g* for 2-4 min. Pellets of undigested tissue were discarded and lysates were purified using the MinElute® PCR Purification Kit (QIAGEN®), with modifications to retain short DNA fragments based on [30]. Final elution in EBT buffer (QIAGEN® EB buffer with 0.05% TWEEN) was performed in two steps with an incubation period of 10 minutes at 37 °C before each centrifugation, for a final volume of 44 µL. Concentration of DNA and fragment size distribution in each of the extracts were assessed with a TapeStation 2200 (Agilent, Santa Clara, CA).

For sequencing library preparation, 100 ng of extracted DNA, in a maximum volume of 32 µL, were used as input template. For less concentrated extracts, 32 µL of the purified lysate were used even if they contained less than 100 ng of DNA (to a minimum of 25 ng). Given the chemically damaged nature of the DNA in the historical samples, we followed the BEST library build protocol [31], a blunt-end, single-tube library preparation procedure suitable for degraded DNA. The library adapters used were custom-designed for the BGISEQ 500 Sequencing Platform [32]. Finished libraries were purified with MinElute® columns (QIAGEN®) and each was eluted in EBT buffer (QIAGEN® EB buffer with 0.05% TWEEN) in two steps, with an incubation period of 10 minutes at 37 °C before each centrifugation, for a final volume of 44 µL. Unamplified libraries were characterised in a qPCR reaction to estimate the number of cycles needed to reach the amplification plateau while avoiding excessive PCR cycling [33].

Libraries were amplified in two reactions of 50 μL each, containing 5 U AmpliTaq Gold polymerase (Applied Biosystems, Foster City, CA), 1× AmpliTaqGold buffer, 2.5 mM MgCl2, 0.4 mg/mL bovine serum albumin (BSA), 0.2 mM each dNTP, 0.2 μM BGI forward primer [32], 0.2 μM BGI reverse index-primer [32] and 10 μL of library DNA template. PCR products were purified with either MinElute® columns (QIAGEN®), or SPRI beads following a double size-selection protocol to retain fragments 100-800 bp (bead-to-DNA ratio of 0.6 in the first selection step, and 1.8 in the second). DNA was eluted in a final volume of 44 μL of EBT buffer with a 10-minute incubation step at 37°C. DNA concentration was measured with a TapeStation 2200 (Agilent, Santa Clara, CA). Amplified libraries that contained >150 ng of DNA were submitted for one lane each of BGISEQ 500 SE100 sequencing.

The nine modern samples were either the fraction of white cells and platelets from blood samples, or pellets of cell cultures. DNA was extracted with phenol-chloroform separation of aqueous and hydrophobic phases. Sequencing libraries were built using the Illumina® TruSeq PCR-free method on DNA inserts that were 350 bp in average fragment length and following manufacturer’s guidelines. Libraries were then sequenced on an Illumina® HiSeq X platform, giving 0.5 lanes per sample in PE150 mode.

### Bioinformatic data processing and quality assessment

#### Quality control, trimming and mapping of DNA sequence data

Raw data included 33 newly re-sequenced genomes from museum specimens, 9 newly re-sequenced genomes from modern samples plus 13 genomes that had been previously published in [18] (NCBI project code: PRJNA394025). Basic quality summaries of the raw data per sample (files in. *fastq* format) were obtained with fastqc v0.11.7 [34]. Then the pipeline PALEOMIX 1.2.13.2 [35] was run for each sample. It included: removal of adapters with AdapterRemoval v2.2.2 [36], alignment against the GCA_000283155.1 white rhinoceros reference genome (NCBI accession code <u>PRJNA74583</u>) [37] with the BWA v0.7.16a *backtrack* algorithm [38], duplicate filtering with Picard MarkDuplicates [39], and assessment of ancient DNA damage with mapDamage v2.0.6 [40]. Minimum base quality filtering was set to zero in order to maximise the number of reads aligned, and customise filtering in later analysis steps.

#### Variant site finding

##### Selection of scaffolds

To optimise memory usage in subsequent analyses, we restricted statistical analysis to scaffolds >13 Mbp in the white rhino reference assembly. The 66 scaffolds above this cut-off were subjected to an analysis of normalised depth of coverage across female and male samples. Three scaffolds showed half the average depth in samples originating from males, a sign that they belong to the X chromosome. These scaffolds were therefore discarded from further analyses. The final set included 63 scaffolds that represent 76.17% of the total assembly.

##### Genotype likelihoods

We identified putative SNPs and calculated genotype likelihoods using ANGSD v 0.921 [41] for the chosen 63 scaffolds, excluding low-complexity regions (a bed file comprising all non-masked regions was provided via the -sites option to ANGSD). Different panels of genotype likelihoods were computed for several sets of samples: for the entire final dataset (n = 52), for a set of non-related samples (n = 49), for each subspecies separately (n_NWR_ = 25; n_SWR_ = 27). In every case, the minimum number of individuals in which a variant site must be present (-minInd) was 95%. Minimum and maximum global depth per site were based on a global depth assessment with ANGSD (-doDepth): 200 and 900 for panels combining both subspecies, 100 and 500 for subspecies-specific panels. Additionally, these quality filtering and output choice parameters were applied: -remove_bads 1 -uniqueOnly 1 -baq 1 -C 50 -minMapQ 30 -minQ 20 -doCounts 1 -GL 1 -doGlf 2 -doMajorMinor 1 -doMaf 1 -doHWE 1 -dosnpstat 1 -HWE_pval 1e-2 -SNP_pval 1e-6 Transitions were removed *a posteriori* with custom-made code.

### Statistical analyses of genomic data

#### Relatedness test

We ran an analysis of relatedness based on a panel of genotype likelihoods with ngsRelate v2 [54], following the approach described in [55]. Two samples appeared as identical (SD1905.5 and SD1905.6) (Figure S3A-B), therefore SD1905.6 was discarded from further analyses. In a separate analysis per subspecies, we found that two pairs of NWR and one pair of SWR samples showed a relatedness signal (Figure S3C-F), so for analyses of structure (i.e. PCA, UMAP and admixture), the sample of lowest depth of coverage from each pair was excluded (CD-un.1, un1856.1, ZA1842.1).

#### PCA analysis and dimensionality reduction with UMAP

To run a principal component analysis, we used PCangsd v 0.973 [42] and a genotype likelihoods panel of transversion sites (n = 818,731 sites) for 49 unrelated samples. The output was a covariance matrix. We then calculated the eigenvectors and eigenvalues using standard packages in R v 3.4.4 [43].

In addition, we reduced the dimensionality of the same panel of genotype likelihoods for n = 49 unrelated individuals with UMAP [22], a non-linear method with which we captured more of the variability in the data and obtained a higher resolution picture of the structure in the dataset. The genotype likelihoods corresponding to 818,731 variable transversion sites were converted into an array and provided as input to the UMAP function *fit_transform*, with a value of nine for the parameter of nearest neighbors. We ran UMAP in python, with modified command lines sourced from the UMAP online tutorial on the iris dataset [44]. The resulting embedding (two-dimensional output) was visualised with *ggplot2* [45] *in R v 3.4.4 [43].*

#### Admixture

Assessment of admixture proportions in our dataset was done by means of ngsAdmix v 32 [46]. Genotype likelihoods, including only transversion sites (n = 818,731 sites) for 49 unrelated samples, were given as input. For each value of K between 2 and 6, 50 runs of ngsAdmix were launched. For each value of K, the run of highest likelihood was chosen for visualization with the specialised software pong [47].

#### Pairwise diversity as π_tv_

For each subspecies separately, a haploid consensus panel of variant sites was computed with ANGSD [41] -doHaploCall. Transitions were omitted, only sites present across 95% of samples were retained (-minInd), and the following quality filtering and output parameters were used: -remove_bads 1 -uniqueOnly 1 -baq 1 -C 50 -minMapQ 30 -minQ 20 -minIndDepth 5 -doCounts 1 -GL 1 -doMajorminor 1. We removed the major allele column from output files, and then used them as input for *popgenWindows.py* [48]. Window size was big enough to encompass entire scaffolds, and the ‘populations’ to compare were the pre-versus post-bottleneck samples. Proportion of pairwise differences within each ‘population’ was visualised in R v 3.4.4 [43].

#### Genome-wide heterozygosity

To assess individual levels of heterozygosity, the folded site-frequency spectrum (SFS) was computed for each sample separately from its corresponding alignment file (.*bam*) with the ANGSD [41] option -dosaf 1 and then with realSFS as indicated in the ANGSD manual [49]. Only the 63 chosen scaffolds were considered (see *Variant site finding*). The following quality filtering parameters were used: -remove_bads 1 -uniqueOnly 1 -baq 1 -C 50 -minMapQ 30 -minQ 20. Since we used the -fold 1 option (we cannot polarise ancestral vs derived alleles), the ancestral (-anc) and the reference (-ref) flags were both fed the white rhinoceros reference assembly.

With the output from ANGSD -dosaf 1 we estimated the SFS with realSFS in chunks of 1 ×10^8^ sites for transversions only (-noTrans 1). The resulting estimates of homozygous and heterozygous sites per chunk were summed up, and the heterozygous sites were divided by the total number of sites to obtain a genome-wide estimate of individual heterozygosity at transversions.

#### Heterozygosity correction

If mean depth of coverage and amount of missing data vary among samples, this could be leading to an inflation or an underestimation of heterozygosity estimates even if using genotype likelihoods. To counteract this, we applied a correction by depth of coverage to estimates of genome-wide heterozygosity.

Mapped reads (.*bam* files) for samples with a mean depth of coverage >15× were randomly subsampled to the equivalent of the following depths: 15×, 12×, 9×, 6× and 3× using SAMtools v 1.9 [50,51]. For each downsampled file we estimated heterozygosity following the procedure described in *Genome-wide heterozygosity*. For each sample, downsampled estimates of heterozygosity were divided by the estimate at the original depth. In this way we obtained the proportion of the maximum/ original heterozygosity estimate at each of the decreasing values of depth of coverage. These proportions were then plotted against the depth of coverage, and we observed that estimates of heterozygosity decreased with depth in a non-linear fashion.

The best fitting curve (a polynomial function of degree three) to this distribution was used to correct the genome-wide heterozygosity estimates of all samples: *gw-heterozygosity = -0.388 + 0.202 * x - 0.01 * x*^*2*^ *+ 1.7E-04 * x*^*3*^, where *x* was replaced by the corresponding value of depth of coverage. The samples of lowest depth of coverage (<3×) appear as outliers in the distribution of heterozygosity despite this correction, reflecting the limitation in sensitivity of this approach at low depths of coverage.

#### Heterozygosity in exons

The very same approach used for the *Genome-wide heterozygosity* was followed, but in this case. *saf* files were generated with ANGSD [41] -dosaf 1 only for those regions falling under the ‘exon’ category along the 63 chosen scaffolds (see *Variant site finding*) according to the publicly available genomic annotation of the white rhinoceros [37]. Once again, only transversion sites were considered. A single, folded SFS for all exonic regions was computed per sample. Heterozygosity was calculated as the ratio between the number of heterozygous sites and the total number of sites in the SFS.

To account for the variability in mean depth of coverage across samples, estimates of heterozygosity in exons were corrected by depth following the same procedure as described in *Heterozygosity correction* and the following polynomial equation: *exon-heterozygosity = -0.35 + 0.188 * x - 8.83E-3 * x*^*2*^ *+ 1.42E-04 * x*^*3*^, where *x* was replaced by the corresponding value of depth of coverage per sample.

#### Heterozygosity in sliding windows and inbreeding measured as F_RoH_

To calculate the proportion of the genome putatively in runs of homozygosity (RoH), we computed local estimates of heterozygosity following the approach described in *Genome-wide heterozygosity*, but providing a list of 3,659 sliding windows of 1 Mbp in length and 0.5 Mbp slide along the 63 chosen scaffolds (see *Variant site finding*). These were used as regions of interest for realSFS (using the parameter flag -r). Input files were the same. *saf* files used for the estimation of genome-wide heterozygosity. Resulting local estimates were not corrected by depth of coverage due to the large burden of computational time and memory it would require.

We visualised the distribution of local estimates of heterozygosity per sample (Figure S7). Based on this, we set an arbitrary threshold of heterozygosity at 5 × 10^−5^ below which heterozygosity is considered low enough to consider that region as part of a RoH.

With a custom script (see *findRoH_v3.py* in [52]) we categorised windows in a binary manner: for each sample, and one scaffold at a time (of a total 63 scaffolds >13 Mbp), windows below the chosen threshold were assigned a value of ‘1’, whereas remaining windows were assigned a ‘0’. The output of this script was a list of regions of low heterozygosity of different lengths, which was represented by the sum of adjacent windows with a heterozygosity value below the threshold (i.e. adjacent windows indexed with a ‘1’). A RoH was declared for each of those regions if n_windows_ ≥ 2. The length of RoHs for each sample was calculated in R v 3.4.4 in the following manner: *RoH_length = n*_*windows*_ *\* 1 Mbp - ((n*_*windows*_ *-1) * 0.5 Mbp)*. With a window size of 1 Mbp and a slide size of 0.5 Mbp, the minimum RoH size considered was 1.5 Mbp (i.e. two adjacent windows).

To obtain estimates of the inbreeding coefficient, we calculated F_RoH_ as the ratio between the length (in base pairs) falling within the declared RoH and the total length scanned, i.e. *total_length = n*_*total_windows*_ *\* 1 Mbp - ((n*_*total_windows*_ *-1) * 0.5 Mbp).*

#### Visualization of genomic diversity metrics

All plots related to genomic erosion metrics were made with *ggplot2* [45] *in R v 3.4.4 [43].*

#### Estimation of N_e_ with demographic modelling

To simulate the recent known demographic histories of NWR and SWR, we computed the 2-dimensional SFS (2d SFS) between the pre- and the post-bottleneck sets of samples for each subspecies separately with ANGSD [41] -doSaf 1 and realSFS. Given the uneven sample sizes between pre- and post-bottleneck, the unfolded 2d SFS was generated first, and then folded with dadi [53], which, unlike ANGSD, can fold asymmetrical 2d SFS. These were then provided as input to fastsimcoal2 v fsc26 [25] under a particular demographic scenario following estimated population sizes and timing of demographic shifts reported in the literature for each subspecies.

The NWR population was simulated under a model of large population size followed by a harsh bottleneck; the SWR demographic model involved the same parameters plus a population expansion after the bottleneck. For each of 5,000 simulation iterations, 50,000 steps were allowed. Mutation rate used was 2.5 × 10^−8^ mutations per generation (the value estimated for humans) as in [18]. The parameters to estimate were time between events in generations, and effective population sizes (in haploid number of chromosomes, later divided by two to obtain the estimated number of individuals).

