## Supplementary material (main) for "Historical population declines prompted significant genomic erosion in the northern and southern white rhinoceros (*Ceratotherium simum*)"

### *Mapping statistics and ancient DNA damage assessment*

Out of a total of 55 white rhinoceros re-sequenced genomes, two were discarded from further analyses because they showed an average depth of coverage <1x. An additional sample was discarded since it was identical to another one sequenced at higher depth of coverage (see *Relatedness test* and Figure S3). Mapping statistics per sample, for the remaining 52, can be found in Table S1. Sample identifiers indicate the code of the country of origin, the date (of collection for pre-bottleneck samples, and of birth for post-bottleneck samples), as well as a counter to distinguish samples of same age and country of origin. Summary statistics of the sequencing data per group are reported in Table S2.

Table S1. *Metadata, mapping statistics and individual diversity measures per sample*. See spreadsheet *WR_TableS1_v3* (available at https://github.com/fasaba/WR_supplementary). The variable *retained_reads* refers to the number of reads that were not discarded by PALEOMIX based on size filtering (>25 bp), and could then be aligned to the reference assembly; the remaining mapping statistics were calculated by PALEOMIX based on this number. The column *depth_of_coverage* refers to the average depth of coverage across the entire reference assembly as calculated by PALEOMIX. The endogenous content of each sample is indicated by *mapped_reads_unique_fraction,* which is the fraction of raw reads mapped minus the fraction of those reads that were clonal.

| | | Retained reads | | Mapped fraction raw | | Clonality | | Mapped fraction unique | | Depth of coverage | | 5'CtoT damage | |
|---|---|---|---|---|---|---|---|---|---|---|---|---|---|
| | n | mean | SD | mean | SD | mean | SD | mean | SD | mean | SD | mean | SD |
| **NWR1** | 16 | 7.73E+08 | 3.22E+07 | 0.7029 | 0.2190 | 0.3132 | 0.1347 | 0.4921 | 0.1979 | 12.10 | 5.42 | 0.0259 | 0.0118 |
| **NWR2** | 9 | 1.80E+08 | 1.34E+07 | 0.7427 | 0.0567 | 0.0039 | 0.0011 | 0.7398 | 0.0562 | 9.25 | 1.05 | 0.0008 | 0.0001 |
| **SWR1** | 9 | 8.04E+08 | 2.56E+07 | 0.6806 | 0.2084 | 0.2939 | 0.0896 | 0.4769 | 0.1629 | 10.88 | 4.66 | 0.0284 | 0.0146 |
| **SWR2** | 18 | 4.41E+08 | 1.87E+08 | 0.7965 | 0.0925 | 0.1430 | 0.0911 | 0.6847 | 0.1103 | 14.92 | 5.15 | 0.0022 | 0.0015 |

Table S2. *Summary statistics of sequence data per group*.

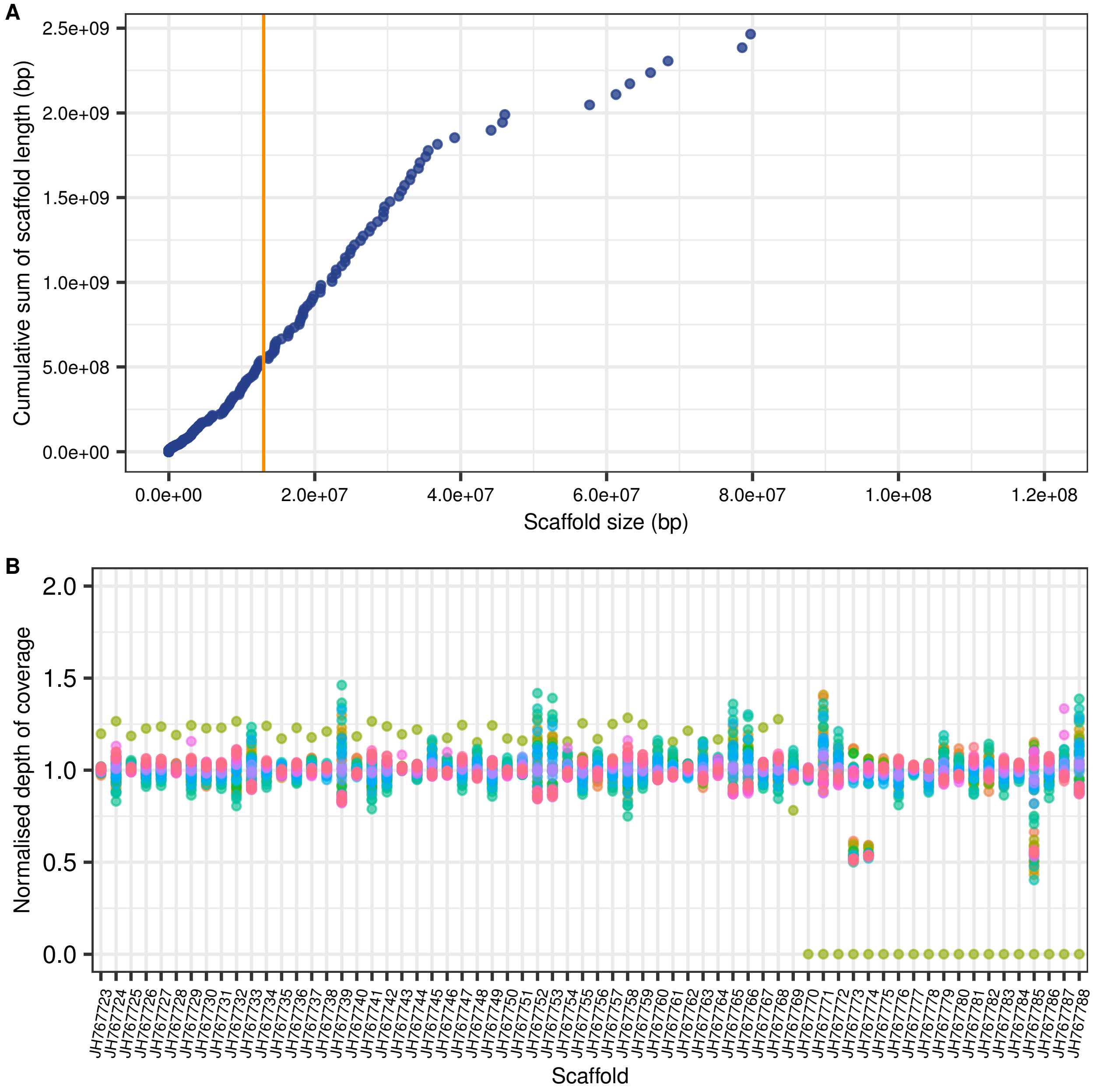


Figure S1. *Choice of scaffolds for variant site finding*. (A) Contribution of each scaffold in the reference assembly to the total length of it. The vertical orange line is placed at a scaffold size of 13 Mbp; only scaffolds above this size were considered for variant site finding. (B) Normalised depth of coverage for each scaffold >13 Mbp for 52 samples. Of the 66 scaffolds on the horizontal axis, three (JH767773, JH767774, JH767785) show a normalised depth of 0.5 for male samples, and were therefore excluded from further analyses.

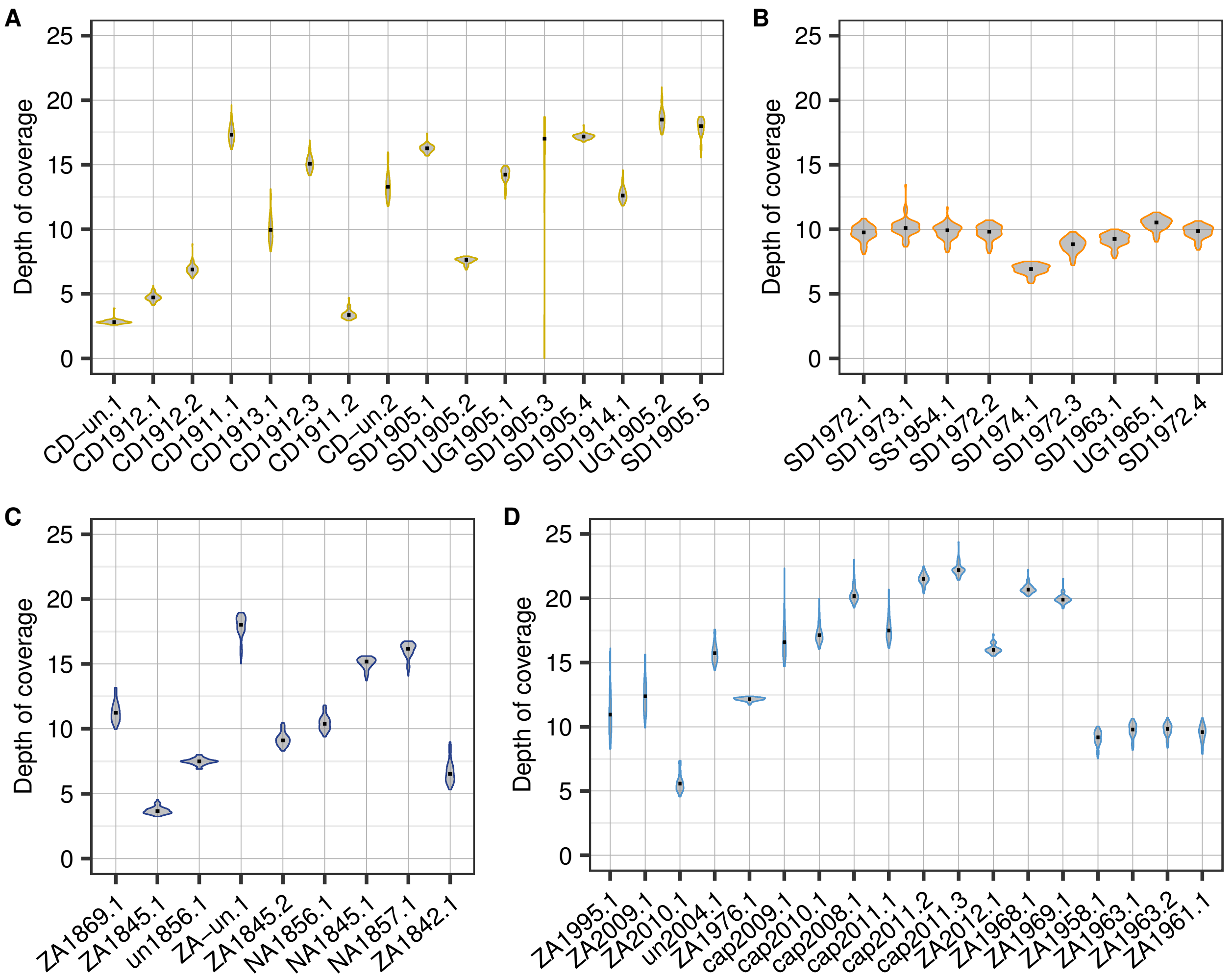


Figure S2. *Distribution of depth of coverage per sample.* For each sample, the distribution of depth of coverage across the 63 chosen scaffolds is shown; samples are sorted into: NWR1 (A), NWR2 (B), SWR1 (C) and SWR1 (D).

#### *Cross-contamination tests*

To verify that no major contamination issues among samples could bias further analyses, we extracted the fraction of reads that mapped against the mitochondrial scaffold (chrM) from each bam file, and generated a fasta file from those reads with Analysis of Next Generation Sequencing Data (ANGSD) v 0.921 [41] option -doFasta 1. The frequency of sites bearing 1, 2, 3, or 4 alleles was computed. Since the mitochondrial chromosome is haploid, for a given sample and site, a frequency of 1 is expected for a given allele; additional alleles occurring might be due to sequencing error, aDNA damage or contamination. Diagnostically, contamination is associated with frequencies above 20% of alternative alleles in these haploid sites; lower frequencies are likely due to errors. We observed that, most samples contained sites with more than one allele in their mtDNA, but at frequencies below 2%.

#### *Relatedness test*

We ran an analysis of relatedness based on a panel of genotype likelihoods with ngsRelate v2 [54], following the approach described in [55]. Two samples appeared as identical (SD1905.5 and SD1905.6) (Figure S3A-B), therefore SD1905.6 was discarded from further analyses. In a separate analysis per subspecies, we found that two pairs of NWR and one pair of SWR samples showed a relatedness signal (Figure S3C-F), so for analyses of structure (i.e. PCA, UMAP and admixture), the sample of lowest depth of coverage from each pair was excluded (CD-un.1, un1856.1, ZA1842.1).

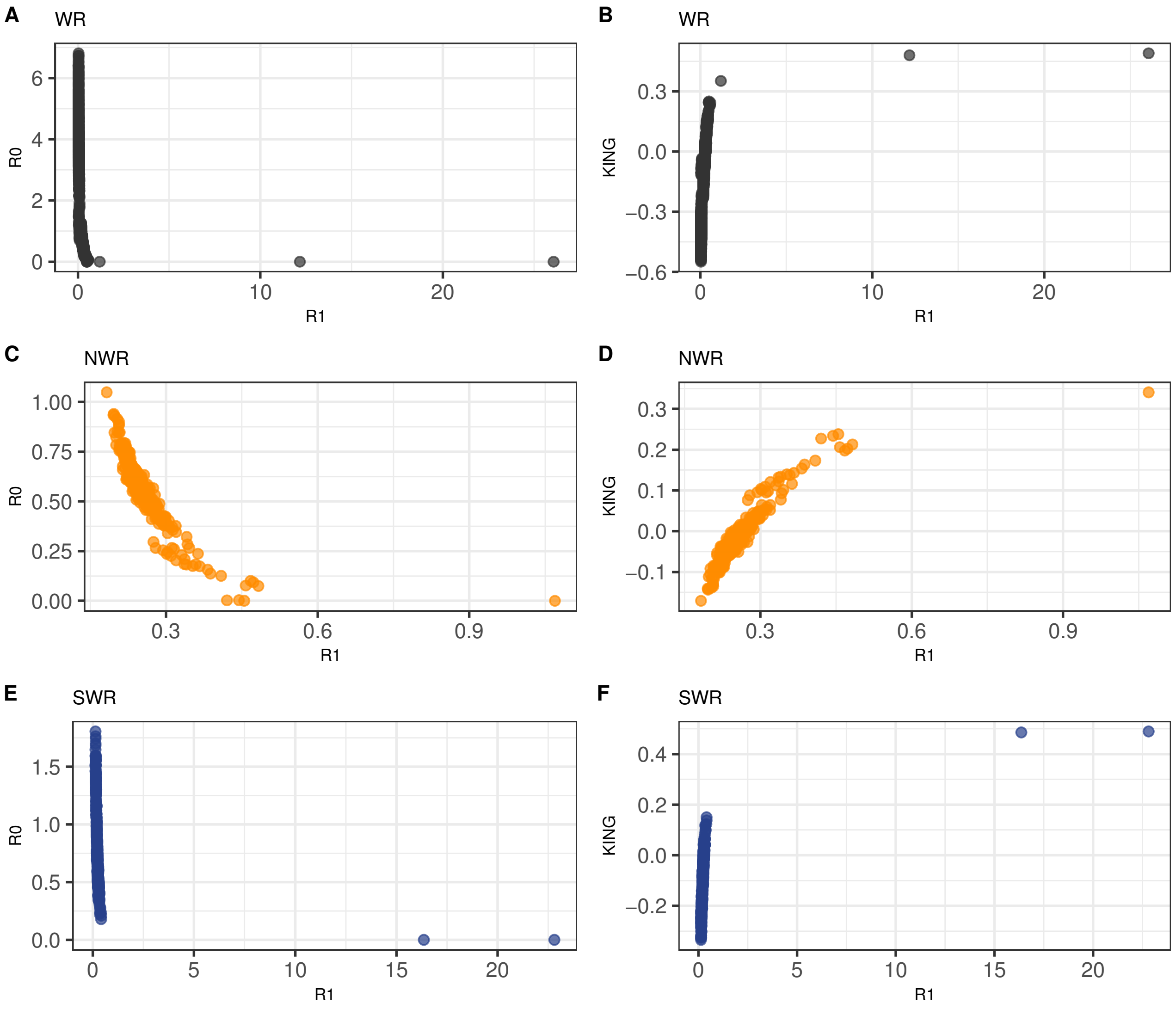


Figure S3. *Relatedness assessment of white rhinoceros samples with ngsRelate*. Analysis of 53 re-sequenced white rhinoceroses (A, B), for 25 NWR (C, D), and for 27 SWR (E, F). In all cases, analyses were based on a panel of genotype likelihoods for transversion sites. For each sample set, each point represents a pairwise combination of samples, and its combination of coefficients is a proxy for the degree of relatedness [55]. In A) and B), the furthest outlier pair corresponds to SD1905.5 - SD1905.6, seemingly identical samples. In C) and D) the related pair corresponds to CD-un.1 - CD-un.2; in E) and F) related pairs are un1856.1 - ZA-un.1 and ZA1845.2 - ZA1842.1.

#### *Principal component analysis (PCA)*

Here are the visualizations of PCs one to three from a PCA analysis of 49 unrelated samples in our dataset. The first principal component (PC1) shows a clear separation between NWR and SWR (Figure S4); the PC2 uncovers population substructure within both subspecies. Post-bottleneck NWR fell within the Sudan-Uganda historical diversity only, but our pre-bottleneck NWR expanded beyond this gradient to represent a wide range of DRC diversity as well. In the SWR, PC2 revealed a split between pre- and post-bottleneck (Figure S4A), where pre-bottleneck samples were more widely scattered, while post-bottleneck SWR clustered tightly together. The PC3 resolved the distribution of SWR, as we observed substructure due to geography in the pre-bottleneck samples, and a comparatively small differentiation among post-bottleneck SWR (Figure S4B).

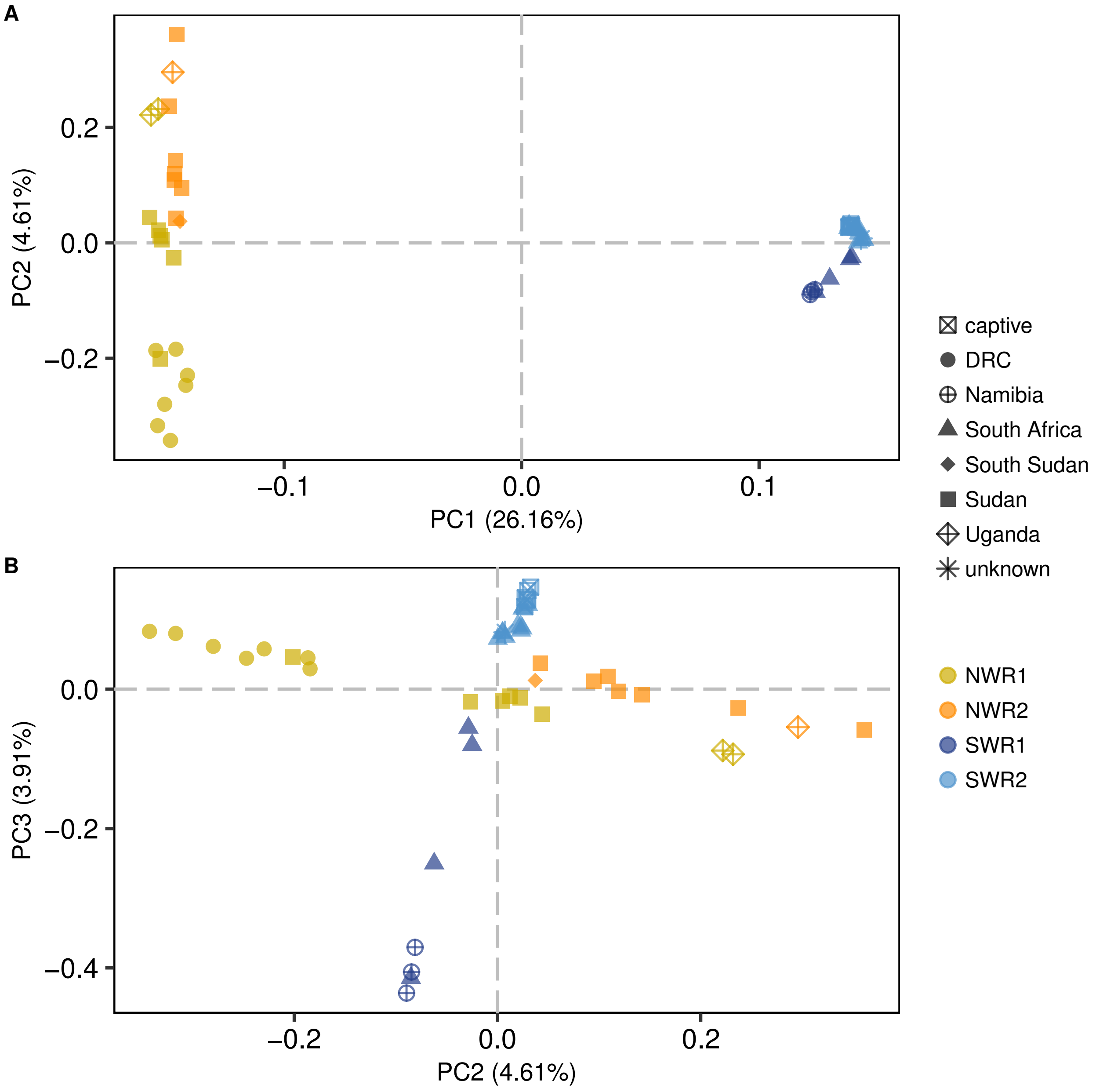


Figure S4. *PCA analysis of 49 unrelated white rhinoceroses*. Based on genotype likelihoods for transversion sites only. A) PC1 against PC2; B) PC2 against PC3.

### ***Heterozygosity correction***

To account for differences in depth of coverage across samples, the estimated genome-wide heterozygosity and exon heterozygosity were corrected based on how heterozygosity estimates decay with decreasing values of depth of coverage (see *Heterozygosity correction* in Methods for more details). The curves from which the correcting equations were drawn are depicted in Figure S5 for genome-wide heterozygosity estimates, and in Figure S6 for exon heterozygosity estimates.

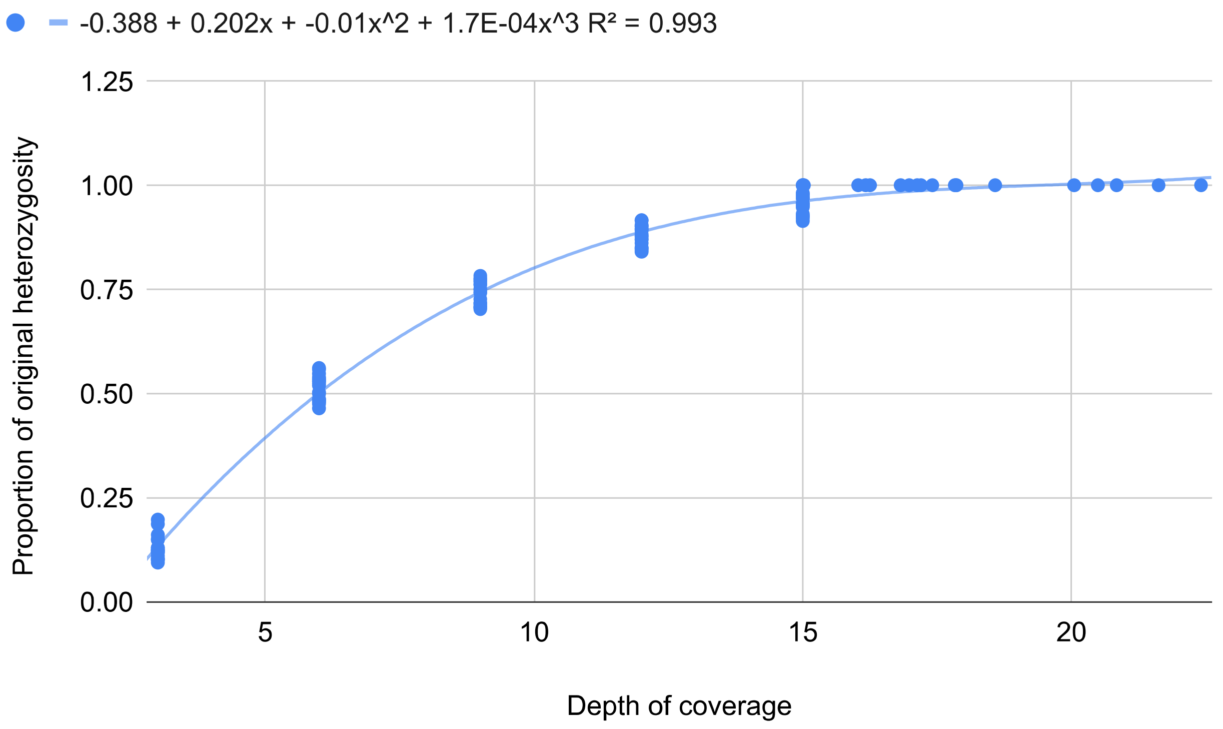


Figure S5. *Relation between the depth of coverage and the normalised genome-wide heterozygosity.* Estimates of genome-wide heterozygosity were calculated based on the individual SFS for decreasing values of depth for samples of mean depth of coverage >15x (n = 19). The trend line was generated in excel by choosing the polynomial equation of lowest degree and highest $R^2$ that best describes the distribution.

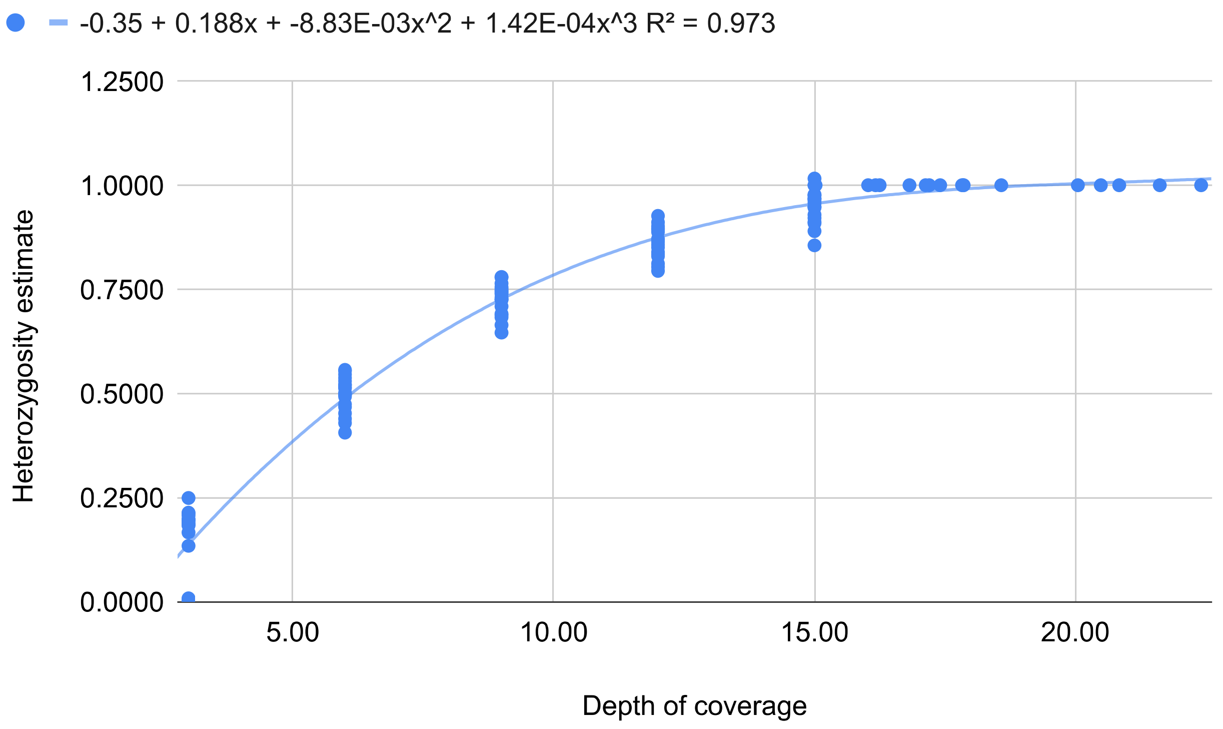


Figure S6. *Relation between the depth of coverage and the normalised exon heterozygosity.* Estimates of exon heterozygosity were calculated based on the individual SFS for decreasing values of depth for samples of mean depth of coverage >15x (n = 19). The trend line was generated in excel by choosing the polynomial equation of lowest degree and highest $R^2$ that best describes the distribution.

### *Local estimates of heterozygosity and identification of RoH*

To calculate an inbreeding coefficient per sample, we visualised the distribution of windows falling along the range of local estimates of heterozygosity per sample (Figure S7). In NWR2 and SWR2, these distributions are skewed toward lower values of heterozygosity. There are three outliers (CD-un.1, CD1911.2 and ZA1845.1) that show a disproportionate number of windows with heterozygosity values very close to zero, probably because they have a substantial amount of missing data given that their mean depth of coverage is <4x. Pre-bottleneck groups, and particularly SWR1, show the longest lengths of RoH detected.

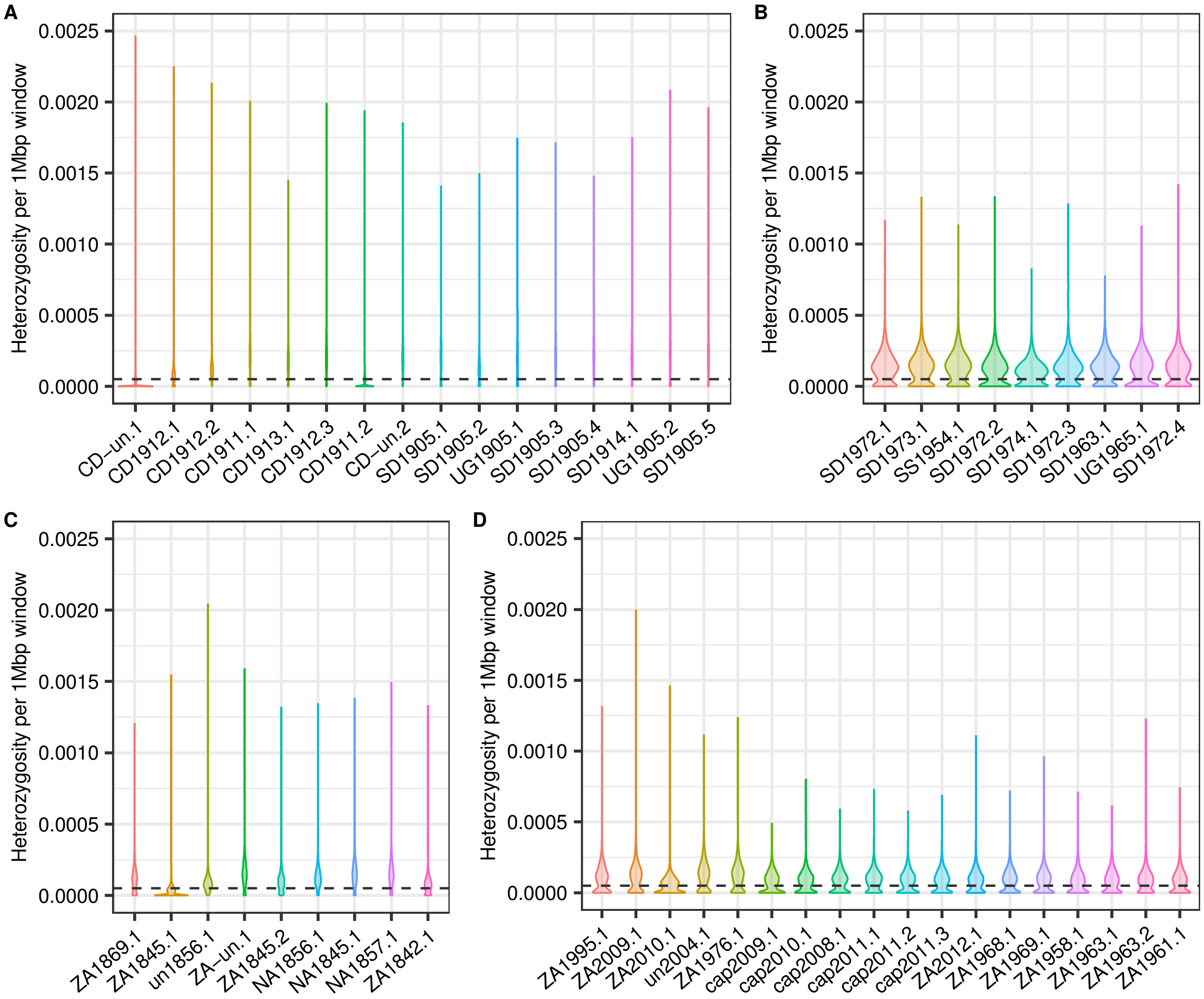


Figure S7. *Distribution of local estimates of heterozygosity per sample and threshold for RoH identification*. Density of windows (1 Mbp, 0.5 Mbp slide) along the range of heterozygosity values per sample for A) NWR1, B) NWR2, C) SWR1 and D) SWR2. Dashed lines indicate the chosen threshold of heterozygosity for assigning windows to RoH ($5 \times 10^{-5}$). Samples CD-un.1 and CD1911.2 (NWR1), and ZA1845.1 (SWR1) were discarded from the $F_{RoH}$ estimation due to an excess of missing data.

#### *Genomic erosion in subsampled post-bottleneck groups*

To exclude that sampling bias might be creating spurious patterns of genomic erosion, we calculated the individual metrics of genomic diversity for a subset of the samples. In NWR1, we kept only samples originating from Sudan-Uganda since all NWR2 were sourced from that area. From SWR2 only wild-born individuals were retained for comparison with all SWR1. This subsampled dataset consists of 38 individuals distributed across the four groups (see Table S3).

Delta estimators for genome-wide heterozygosity, exon heterozygosity and $F_{RoH}$, show that, after minimizing sampling bias, genomic erosion patterns remain (Figures S8 and S9). Post-bottleneck NWR and SWR show significantly lower levels of heterozygosity than pre-bottleneck counterparts (see Figure S8 and Table S3 for delta estimators), with the exception of $F_{RoH}$ between NWR1 and NWR2, despite a slight increase among post-bottleneck NWR (Figure S9 and Table S3). Wild-born SWR2 remain significantly more inbred than SWR1 (Figure S9 and Table S3).

| | n | Date of collection / birth | Median GW het | Unpaired Wilcoxon test p-value | Δ GW het | Median exon het | Unpaired Wilcoxon test p-value | Δ exon het | Median $F_{RoH}$ | Unpaired Wilcoxon test p-value | Δ $F_{RoH}$ |
|---|---|---|---|---|---|---|---|---|---|---|---|
| **NWR1 SD-UG** | 8 | 1905-1914 | 0.000192 | 1.65E-04 | -0.0827 | 0.000112 | 2.74E-02 | -0.1431 | 0.1867 | 1.14E-01 | 0.1426 |
| **NWR2** | 9 | 1954-1974 | 0.000176 | | | 0.000096 | | | 0.2134 | | |
| **SWR1** | 9 | 1905-1914 | 0.000158 | 2.04E-04 | -0.2764 | 0.000087 | 3.39E-02 | -0.1660 | 0.1831 | 5.49E-03 | 0.6761 |
| **SWR2 wild** | 12 | 1954-1974 | 0.000114 | | | 0.000073 | | | 0.3068 | | |

Table S3. *Overview of the subsampled dataset and summary of delta estimators*. For NWR1 and SWR1, time span refers to collection dates; for NWR2 and SWR2, it refers to birth dates. For individual genome-wide heterozygosity, exon heterozygosity and $F_{RoH}$, comparisons of the medians were calculated with Wilcoxon tests. Delta estimators were calculated as median pre-bottleneck value minus median post-bottleneck value, divided by the median pre-bottleneck value. All metrics were based on transversion sites only.

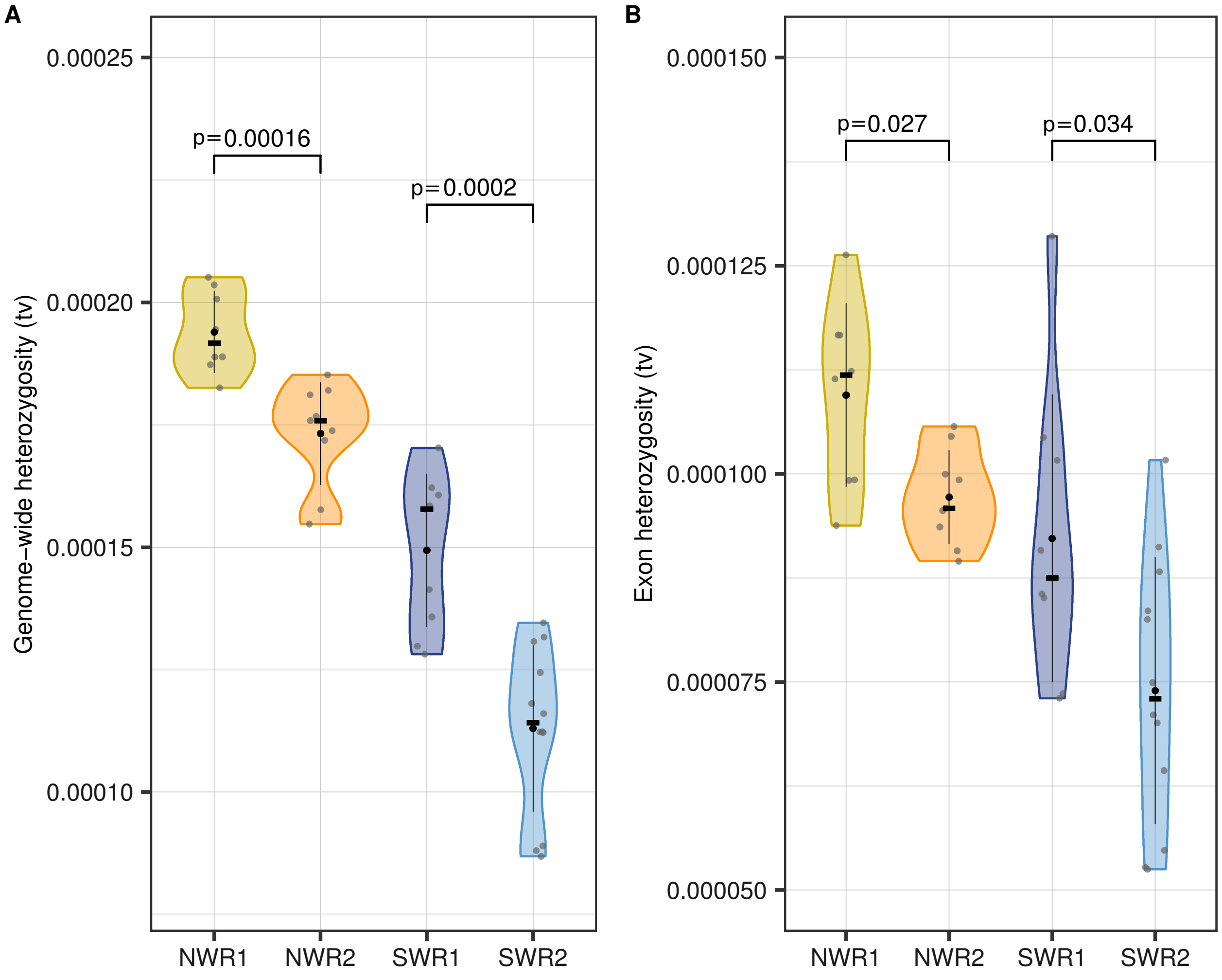


Figure S8. *Post-bottleneck white rhinoceroses show lower genomic diversity after subsampling to minimise sampling bias*. Estimates of genome-wide heterozygosity (A) and heterozygosity at regions annotated as exons (B) based on the per-sample SFS calculated for transversions and then corrected for depth of coverage for the subsampled dataset (n = 38 individuals). Black dots indicate the mean, black lines the standard deviation, and black cross-bars, the median per group. P-values correspond to median comparisons with unpaired Wilcoxon tests.

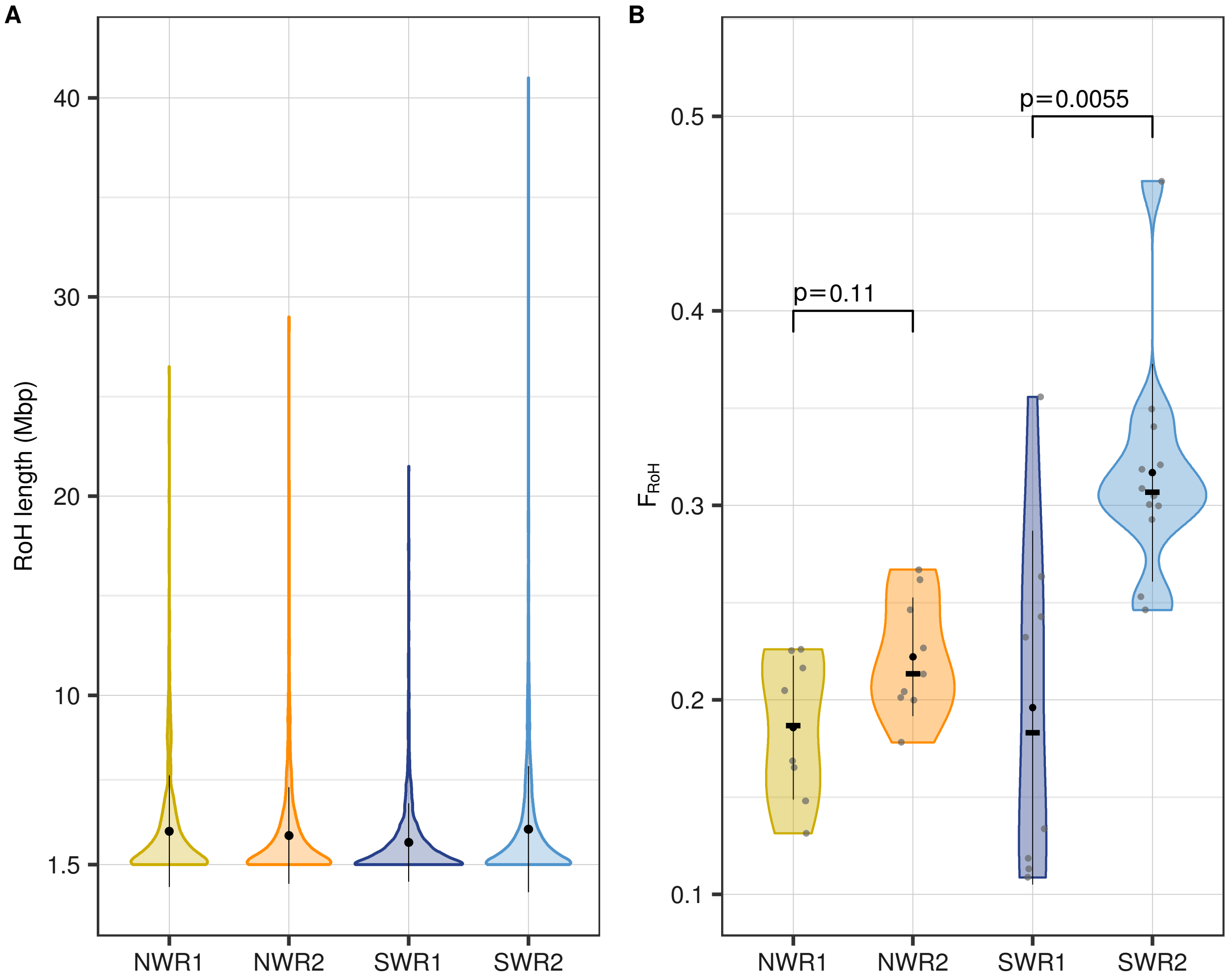


Figure S9. *Post-bottleneck white rhinoceroses show higher estimates of inbreeding after subsampling to minimise sampling bias*. A) Distribution of the length of RoHs for each of the four groups after subsampling and removing one outlier (n = 37 individuals); black dots indicate the mean RoH length per group and black lines the standard deviation. B) Estimates of individual $F_{RoH}$ across groups after subsampling and removing one outlier (n = 37 individuals). Black dots indicate the mean, black lines the standard deviation, and black cross-bars the median per group. P-values refer to unpaired Wilcoxon tests to compare the medians.
